# Systematic Elucidation and Pharmacological Targeting of Tumor-Infiltrating Regulatory T Cell Master Regulators

**DOI:** 10.1101/2022.02.22.481404

**Authors:** Aleksandar Obradovic, Casey Ager, Mikko Turunen, Thomas Nirschl, Mohsen Khosravi-Maharlooei, Christopher Jackson, Vassan Yegnasubramanian, Angelo DeMarzo, Christina Kochel, Mohamad Allaf, Trinity Bivalacqua, Michael Lim, Charles Karan, Charles G. Drake, Andrea Califano

**Affiliations:** Columbia Center for Translational Immunology, Irving Medical Center, New York, NY USA; Department of Systems Biology, Columbia University Irving Medical Center, New York, NY USA; Department of Hematology Oncology, Columbia University Irving Medical Center, New York, NY USA; Department of Oncology, The Johns Hopkins University School of Medicine, Baltimore, MD; Department of Neurosurgery, The Johns Hopkins University School of Medicine, Baltimore, MD; Department of Urology, The Johns Hopkins University School of Medicine, Baltimore, MD; Department of Neurosurgery, Stanford School of Medicine, Palo Alto, CA; J.P. Sulzberger Columbia Genome Center, Columbia University, New York, NY; Herbert Irving Comprehensive Cancer Center, Columbia University Irving Medical Center, New York, NY USA; Janssen Research and Development, Springhouse, PA, USA; Department of Medicine, Columbia University Irving Medical Center, New York, NY USA; Department of Biochemistry & Molecular Biophysics, Columbia University Irving Medical Center, New York, NY USA; Department of Biomedical Informatics, Columbia University Irving Medical Center, New York, NY USA

**Keywords:** Tumor Immunology, T regulatory Cells, Master Regulator Analysis, TRSP1, gemcitabine, Cancer Systems Biology

## Abstract

Due to their immunosuppressive role, tumor-infiltrating regulatory T cells (TI-Tregs) represent attractive therapeutic targets. Analysis of TI vs. peripheral Tregs (P-Tregs) from 36 patients, across four malignancies, identified 17 candidate Master Regulators (MRs), predicted to mechanistically regulate TI-Tregs transcriptional state. *In vivo*, pooled CRISPR-KO screening, using a hematopoietic stem cell transplant model, confirmed essentiality of 7 of 17 MRs in TI-Treg recruitment and/or retention to the TME, without affecting other T cell subtypes, while individual knockout of the most significant MR (TRPS1) significantly reduced tumor allograft growth. TI-Treg drug perturbation profile analysis identified drugs capable of inverting the TI-Treg-specific MR activity signature at low concentration. Low dose treatment with gemcitabine (top prediction) inhibited tumor growth in immunocompetent but not immunocompromised allografts, increased PD-1 inhibitor efficacy, and depleted TI-Tregs *in vivo*. The study provides key insight into Treg infiltration mechanism and a gene reporter assay to identify additional small molecule inhibitors.

**Graphical Abstract:** 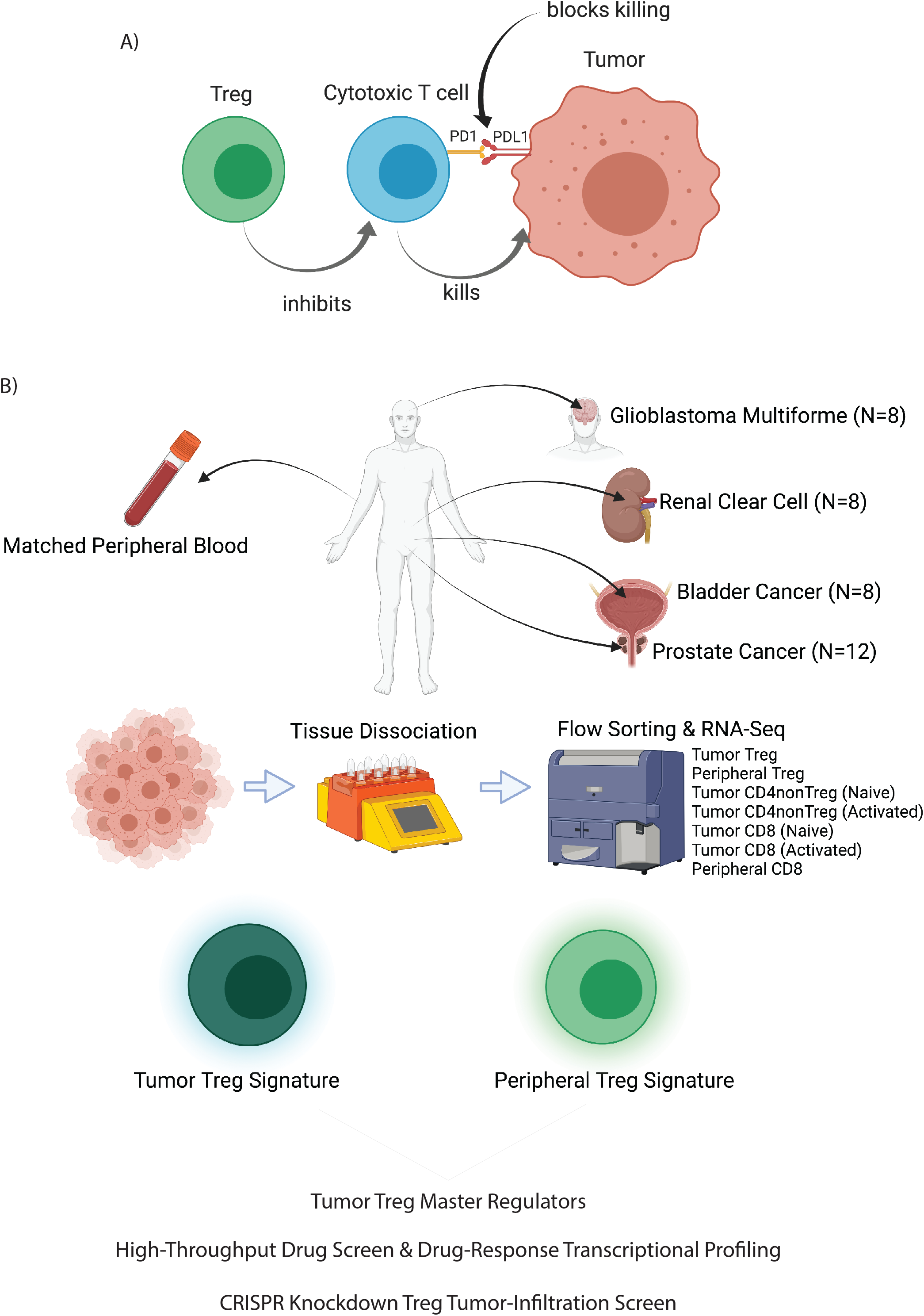

## Introduction

To manifest as clinically apparent disease, cancer must evade a complex repertoire of host-protective immune response mechanisms, the outcome of which is largely determined by the balance of inflammatory (anti-tumor) and tolerogenic (pro-tumor) immune cell function in the tumor microenvironment (TME) (Schreiber et al., 2011). By contributing to a tolerogenic TME, the regulatory T cell (Treg) lineage—characterized by activation of the hallmark transcription factor FoxP3—promotes tumor growth and immunotherapy resistance. As such, increased Treg infiltration in the TME correlates with poor prognosis and increased resistance to immune checkpoint inhibitors across many human malignancies (Chao and Savage, 2018; Flammiger et al., 2013; Muroyama et al., 2017; Obradovic et al., 2020; Shang et al., 2015). While this makes Tregs attractive therapeutic targets, several factors have prevented clinical translation of this concept. First, to avoid severe autoimmunity-mediated toxicity (Bos et al., 2013; Chao and Savage, 2018), an optimal Treg-directed therapy should target tumor infiltrating Tregs (TI-Tregs) while sparing peripheral Tregs (P-Tregs). Second, the Treg transcriptional profile broadly recapitulates that of other activated T cells, thus complicating design of selective targeting strategies that would preserve anti-tumor cytotoxic CD8 and CD4 T cell function (Chao and Savage, 2018; Freeman et al., 2020). The majority of current Treg-targeting agents do not satisfy these criteria and, although effective in murine models, they have not effectively translated to human patients (Selby et al., 2013; Sharma et al., 2019; Simpson et al., 2013). This highlights the need for elucidating the still elusive causal mechanisms that underlie Treg recruitment, retention, and/or function in the TME, thus leading to identification of more specific TI-Treg vulnerabilities.

To address this challenge several labs have profiled Tregs isolated from clinical tumor specimens to identify genes differentially expressed in TI vs. non-TI-Tregs and other T cells. However, differences identified so far have failed to provide mechanisms of tumor infiltration that yield successful pharmacologic targets. For instance, several studies have successfully validated known TI-Treg biology, including high expression of *IL2RA* (CD25) and *FOXP3* in conjunction with multiple T cell checkpoints (*CTLA4, PDCD1* [PD-1]*, HAVCR2* [LAG-3]*, TIGIT*), TNF-family receptors (*TNFSF9* [4-1BB], *TNFRSF18* [GITR], *TNFRSF4* [OX-40]), wound-healing factors (*ENTPD1* [CD39], *IL1RL1* [ST2]), and proliferation programs (De Simone et al., 2016; Magnuson et al., 2018; Plitas et al., 2016; Zheng et al., 2017). In addition, a number of specific genes enriched in TI-Tregs have been observed across datasets, including *LAYN*, *SAMSN1*, *IL1R2*, *MAGEH1, CD177*, and the chemokine receptor *CCR8*. Of these, CCR8 is one leading candidate, showing preferential protein level expression in breast cancer (Plitas et al., 2016) and NSCLC (Van Damme et al., 2021) TI-Tregs, among others. Monoclonal antibodies targeting CCR8 are currently in early-phase clinical trials. However, more recent data suggest CCR8 may be dispensable for Treg function (Van Damme et al., 2021; Whiteside et al., 2021). Thus, despite these advancements, additional efforts are warranted to discover novel potential TI-Treg vulnerabilities via orthogonal approaches.

We have developed methodologies for the assembly and interrogation of lineage context-specific gene regulatory networks, including the Algorithm for the Reconstruction of Accurate Cellular Networks (ARACNe) (Basso et al., 2005) and the Virtual Proteomics by Enriched Regulon analysis (VIPER) algorithm (Alvarez et al., 2016), respectively. These have been successful in nominating Master Regulator (MR) proteins, representing mechanistic drivers of both pathophysiologic and transformed transcriptional cell state (Aytes et al., 2014; Carro et al., 2010; Paull et al., 2021), that have been experimentally validated, including at the single cell level (Obradovic et al., 2021b; Son et al., 2021), see (Califano and Alvarez, 2017) for a recent perspective. Here, we sought to leverage these methodologies to interrogate a Treg-specific gene regulatory network with signatures of TI vs. P-Tregs, to first identify and then validate novel causal Master Regulators presiding over Treg infiltration and retainment to the TME.

To generate tumor-agnostic signatures of TI-Treg vs. P-Treg state, we first assembled patient-matched transcriptional profiles from multiple T cell subpopulations, flow sorted from the tumors and peripheral blood of 36 patients, using established antibody panels. For this study, we focused on tumor types whose T cell repertoire is not well-represented in existing datasets, including prostate adenocarcinoma, bladder cancer, clear cell renal carcinoma, and glioblastoma. This dataset uniquely facilitates identification of transcriptional signatures differentially expressed in TI-Tregs from highly diverse cancers, compared to peripheral blood Tregs, conventional non-Treg CD4 (Tconv), and CD8 T cells. Additionally, it allowed accurate identification of genes specifically induced upon T cell activation, many of which are shared between TI-Tregs and other activated T cell lineages.

Akin to a highly multiplexed gene reporter assay, we then used VIPER to identify MR proteins whose transcriptional targets were most differentially expressed in TI-vs. P-Tregs, thus representing the protein subset (*module*) most likely to mechanistically implement and homeostatically maintain the TI-Treg transcriptional state (Alvarez et al., 2016). We have shown that this approach significantly outperforms gene expression based analyses and compares favorably with single cell, antibody-based approaches, as shown by analysis of single cells RNA-seq profiles (scRNA-seq) (Elyada et al., 2019; Obradovic et al., 2021a).

The analysis, followed by Random Forest-based protein selection (see methods), identified 17 highly significant candidate MRs (*p* ≤ 0.01, Bonferroni-corrected), predicted to physically control the TI- vs. P-Treg transcriptional signature. Most notably, none of the genes encoding for these MRs were significantly differentially expressed or reported in previous studies. This is not unexpected, since these proteins are likely to be post-translationally rather than transcriptionally activated—*i.e.*, by signaling pathways activated by chemokines and other tumor-secreted signals. Notably, VIPER-inferred MRs were virtually identical across four different tumors (Table 1), suggesting that tumor infiltration mechanisms are largely tumor agnostic.

**Table 1:** GSEA of TI vs. P-Treg MR proteins between tumor contexts, Bonferroni corrected.

| Candidate MRs | Differentially Active Proteins | P-value | Candidate MRs | Differentially Active Proteins | P-value |
| --- | --- | --- | --- | --- | --- |
| GBM | PRAD | 4.70E-07 | PRAD | GBM | 1.30E-11 |
| GBM | KIRC | 7.20E-07 | KIRC | GBM | 6.90E-11 |
| GBM | BLCA | 2.10E-05 | BLCA | GBM | 1.10E-06 |
| PRAD | KIRC | 3.40E-11 | KIRC | PRAD | 2.20E-04 |
| PRAD | BLCA | 8.10E-08 | BLCA | PRAD | 2.30E-07 |
| KIRC | BLCA | 2.10E-05 | BLCA | KIRC | 1.30E-08 |

To assess whether candidate MR proteins were essential for TI-Treg infiltration and retention to the TME, we leveraged two orthogonal yet complementary methodologies. First, we utilized a CHIME (CHimeric IMmune Editing (LaFleur et al., 2019)) model to perform a pooled, *in vivo* CRISPR/Cas9 screen to assess whether knockout of candidate MRs would deplete TI-Tregs without affective P-Tregs, thus confirming their mechanistic role in mediating naïve Treg recruitment and/or TI-Treg retention to the tumor microenvironment. Second, we performed a systematic drug screen where patient-derived TI-Tregs were expanded *ex vivo* and then RNA-seq profiled, following treatment with a large compound library. Candidate MR-inverter compounds—capable of specifically inverting the activity of the TI-Treg MRs—were nominated using the NY/CA Dept. of Health approved, CLIA-compliant OncoTreat algorithm (Alvarez et al., 2018). Finally, we validated the ability of the most significant drug (gemcitabine) to phenocopy the MR knockout effects by assessing the effect of gemcitabine on tumor growth rate in immunocompetent versus immunodeficient animals and in conjunction with anti-PD-1 checkpoint blockade immunotherapy.

Taken together, these results show MR analysis is effective in elucidating causal determinants of human TI-Treg transcriptional state and in nominating mechanism-based, clinically actionable therapeutic strategies for modulating TI-Treg infiltration and retention to the TME, thus potentiating immunotherapy. A critical value of this approach is its highly generalizable nature, which permits relatively trivial extension to nominate MRs and MR-inverter drugs for a variety of additional tumor-promoting immune subpopulations in the TME..

## Results

### Isolating tumor (TI-Tregs) vs. blood derived (P-Tregs) Regulatory T Cells

Tumor and patient-matched peripheral blood tissues were collected from 36 individuals, including 8 glioblastoma (GBM), 8 bladder adenocarcinoma (BLCA), 8 clear cell renal carcinoma (KIRC), and 12 prostate adenocarcinoma (PRAD) patients. Multiple T cell lineages were freshly sorted from each patient by antibody-based flow cytometry, including TI-Tregs, P-Tregs, peripheral blood CD4 T cells, and both tumor-infiltrating and peripheral blood CD8 T cells, see Figure S1 for sorting strategies. Purity was assessed by flow and reported to exceed 95% for each population (Figure S1). To provide additional controls for T cell activation, patient-matched aliquots of peripheral (*i.e.*, blood-derived) CD4 and CD8 T cells from each of the 36 patients were stimulated for 24-hours with anti-CD3/anti-CD28 beads. Total RNA was purified from each of these seven distinct T cell subpopulations and RNA-seq profiles were generated by Illumina sequencing, for a total of 252 distinct RNA-seq profiles.

### Nominating Candidate Master Regulators of TI-Treg Transcriptional State

Gene expression-based cluster analysis produced poor stratification of different T-cell subtypes (Figure 1A). Having shown that protein activity-based cluster analysis, using the regulatory network-based VIPER algorithm (Alvarez et al., 2016), consistently outperforms expression-based analyses (Obradovic et al., 2021a; Paull et al., 2021), we proceeded to generate a tumor-infiltrating regulatory T cell-specific gene regulatory network by analyzing the 236 T cell-derived profiles using AP-ARACNe (Lachmann et al., 2016)—the latest incarnation of the ARACNe algorithm (Basso et al., 2005)—followed by VIPER-based measurement of differential protein activity for each sample against the average of all samples, as previously described in multiple publications (Alvarez et al., 2016), see methods.

**Figure 1:**
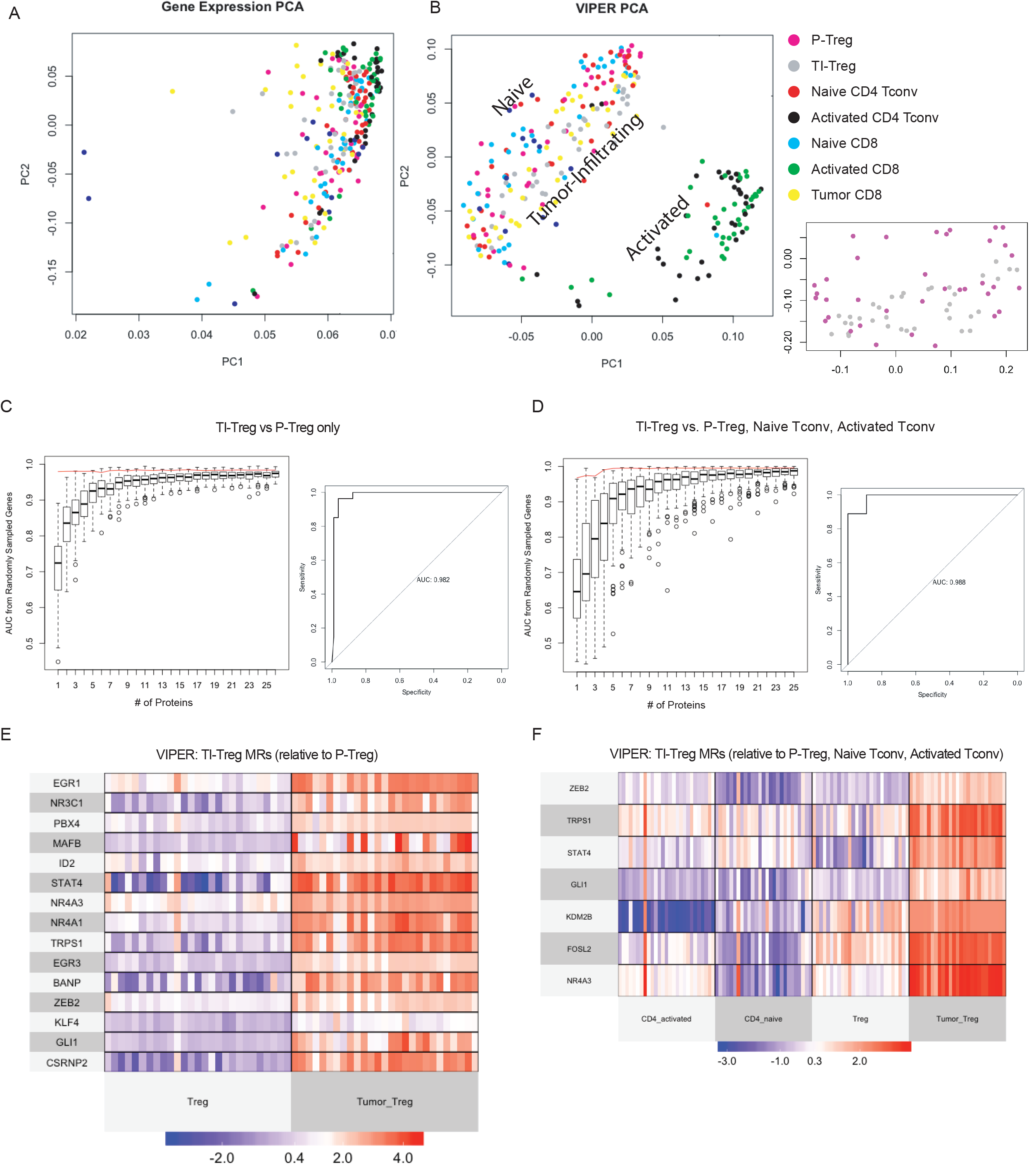
VIPER Enables Definition of Tumor vs Peripheral Treg Master Regulator Signature: **(A)** Principal Component Analysis (PCA) plot of Gene Expression colored by T-cell subtype where black indicates activated CD4nonTregs, green indicates activated CD8 T-cells, red indicates naïve CD4nonTregs, cyan indicated naïve CD8 T-cells, yellow indicates Tumor CD8 T-cells, purple indicates Peripheral Tregs, and grey indicates Tumor Tregs. **(B)** PCA plot of VIPER-inferred protein activity, colored as in A, showing spatial separation of T-cell sub-types. Plot to the lower right showing PCA of VIPER-inferred protein activity separating TI-Tregs and P-Tregs only. **(C)** Random Forest Feature Selection of VIPER Master Regulators Upregulated in Tumor Tregs vs Peripheral Tregs, normalized against Naïve CD4nonTregs. Boxplot shows distribution of test-AUCs for randomly sampled number of genes corresponding to x-axis, with red line indicating actual AUC of selected Master Regulator gene set. AUC of master regulator gene set for selected number of AUCs is shown in inset to the right. **(D)** Random Forest Feature Selection of VIPER Master Regulators Up-Regulated in Tumor Tregs vs Peripheral Tregs, Naïve CD4nonTregs, and Activated CD4nonTregs, normalized against Naïve CD8 T-cells. Boxplot shows distribution of test-AUCs for randomly sampled number of genes corresponding to x-axis, with red line indicating actual AUC of selected Master Regulator gene set. AUC of master regulator gene set for selected number of AUCs is shown in inset to the right. **(E)** Heatmap of VIPER Protein Activity for Master Regulators Selected in C. **(F)** Heatmap of VIPER Protein Activity for Master Regulators Selected in D.

Activity-based cluster analysis showed clear separation of naïve and activated cells by 2D principal component analysis, with tumor-infiltrating cells a distinct cluster (Figure 1B). Intriguingly, neither gene expression nor protein activity could stratify TI-Treg samples by tumor type, suggesting a relatively tumor-agnostic transcriptional state. Indeed, a Random Forest classifier for TI- vs. P-Treg state, independently trained on samples from each tumor type (*e.g.*, using GBM samples only), could perfectly classify TI- vs. P-Tregs across all other tumor types (*i.e.*, PRAD, BLCA, and KIRC) (pairwise Area Under the Receiver Operating Curve, AUROC = 1.0 for all comparisons). Consistently, there was highly significant enrichment (ranging from p = 10^-4^ to p = 10^-11^) of candidate MRs inferred from only one tumor type— based on VIPER analysis of genes differentially expressed in tumor patient-specific TI- vs. P- Tregs (*p* ≤ 10^-3^)—in proteins differentially active in TI- vs. P-Tregs from all other tumor types, by Gene Set Enrichment Analysis (GSEA) (Subramanian et al., 2005) (Table 1).

To select the most discriminative candidate MRs, among those differentially active in TI-Tregs vs. other T cell populations—including P-Tregs, naïve CD4, and activated CD4 T cells—we used the Random Forest algorithm (see methods). Specifically, the analysis selected the minimal number of features (*i.e.*, candidate MRs, starting from the most statistically significant one) that maximized the ratio between the AUROC from a Monte Carlo cross validation (MCCV) analysis, compared to the null hypothesis where features were selected at random from all transcriptional regulators (Figure 1C-D). The analysis yielded 15 proteins significantly differentially active in TI- vs. P-Tregs Figure 1E) (AUROC = 0.982 for TI vs. P-Treg classification by MCCV) (Figure 1C). In addition, seven candidate MRs were found to optimally classify TI-Tregs vs. other control subpopulations, (Figure 1F) (AUROC = 0.988, by MCCV) (Figure 1D). Only two of these seven proteins were not included in the 15 from the TI- vs. P-Treg analysis, yielding a total of 17 unique candidate MRs of Treg tumor infiltration. These include EGR1, NR3C1, PBX4, MAFB, ID2, STAT4, NR4A3, NR4A1, TRPS1, EGR3, BANP, ZEB2, KLF4, GLI1, CSRNP2, KDM2B, and FOSL2. Of these, the NR4A family of transcription factors (Bandukwala and Rao, 2013) as well as FOSL2 (Renoux et al., 2020) were previously reported as upstream regulators of FOXP3 expression in Tregs, the glucocorticoid receptor NR3C1 was shown to have Treg-specific function (Rocamora-Reverte et al., 2019), and EGR3 was reported as a negative regulator of T-cell activation (Morita et al., 2016). However, none were previously reported as causal regulators of Treg tumor infiltration and none was significantly differentially expressed at the RNA level in TI-Tregs in our dataset.

### Candidate MR Validation by In Vivo Pooled CRISPR-KO Screen

To functionally validate whether candidate MRs are essential for TI-Treg recruitment and/or retention to the TME, we performed an *in vivo* pooled CRISPR knockdown screen using the CHIME (CHimeric IMmune Editing) system (LaFleur et al., 2019). Briefly, we sorted Lin^-^Sca-1^+^c-Kit^-^ cells enriched for hematopoietic stem cells (HSCs) from constitutive Cas9-expressing mice, and lentivirally transduced them with a sgRNA library targeting 34 genes, with 3 guides/gene, for a total of 102 guides. Target genes included the 17 MRs described above, 13 randomly selected negative control genes, and 4 positive controls with knockout known to be toxic to Tregs (*Cd4*: toxic to all CD4 T-cells, *Foxp3*: toxic to Tregs, *Plk1* and *Cdk1*: toxic to all cells) (Fig 2A-B). Guides were cloned in the pXPR_053 vector (see methods), with a VexGFP (Vex) fluorophore for transduced cell selection. HSCs were then implanted into irradiated Cas9-tolerized recipients, allowing the immune system to reconstitute *de novo* over 10 weeks, such that all Vex+ immune lineage cells, including Tregs, harbored co-expression of a given guide RNA and Cas9. Syngeneic MC38 colon carcinoma tumors, chosen for their well-established reliance on an intact TI-Treg compartment for *in vivo* growth (Arce Vargas et al., 2018), were implanted and allowed to grow for approximately three weeks. Finally, Vex+ Tregs as well as CD4 Tconv were flow-sorted from the tumor and spleen (control) of each mouse (Fig 2C). The latter was selected as an effective reservoir of P-Tregs—such that differential sgRNA barcode abundance could be compared in TI-Tregs vs spleen P-Tregs.

**Figure 2:**
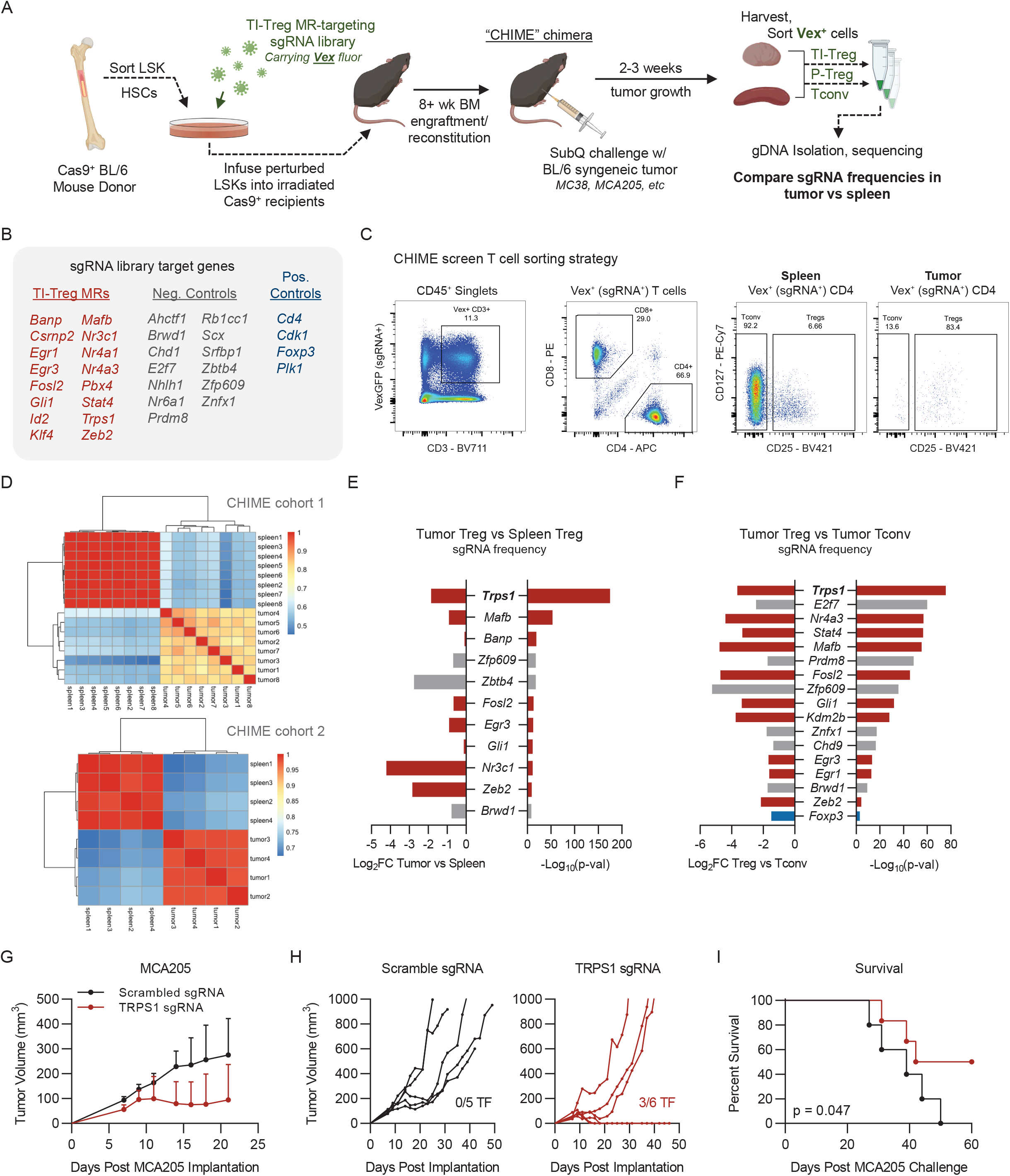
Chimeric Immune Editing Mouse Model Enables Validation of Treg Tumor-Infiltration Master Regulators: **(A)** Experimental design for CRISPRko Validation of Tumor vs Peripheral Treg Master Regulators MR targets, randomly sampled negative control genes, and Treg-toxic positive control genes are listed in the Figure. **(B)** List of sgRNAs targeting 17 TI-Treg MRs, 13 negative control genes, and 4 positive control genes. **(C)** Representative flow cytometry gating for Vex+ CRISPR-transduced Tregs, CD4nonTregs, and CD8 T-cells in spleen and Tumor. **(D)** Correlation of sgDNA frequency distribution between replicates of Spleen and Tumor Tregs in experimental cohorts 1 (top) and 2 (bottom). **(E)** Plots of log2(Fold Change) and Bonferroni-corrected p-values Stouffer-integrated across experimental cohorts for genes with consistent and statistically significant depletion of targeting gDNAs in Tumor Tregs vs Spleen Tregs. **(F)** Table of log2(Fold Change) and Bonferroni-corrected p-values Stouffer-integrated across experimental cohorts for genes with consistent and statistically significant depletion of targeting sgDNAs in Tumor Tregs vs Tumor CD4-nonTregs. **(G)** Tumor growth curves of MCA205 (8×10^5^ implanted subcutaneously) in mice bearing single gene TRPS1 knockdown or scrambled control guide. **(H)** Individual growth curves of mice in (G), with numbers of mice tumor free (TF) noted. **(I)** Kaplan-Meier plot for experiment in (G,H) showing significant difference in tumor growth (p<0.05)

Upon engraftment and reconstitution of the hematopoietic system, roughly 25-40% of immune cells harbored CRISPR gene knockdown (Figure S2A). To assess reproducibility of any observed effects, two separate CHIME chimera cohorts were implanted with syngeneic MC38 tumors, the second being implanted with Lin^-^Sca-1^+^c-Kit^-^ hematopoietic stem cells from the bone marrow of the first. Tumors were grown for 18 days before CD4^+^CD25^+^ Tregs and CD4^+^CD25^-^ Tconv were sorted from the tumor and spleen of each animal (Figure 2C) and sequenced to assess differences in sgRNA representation. Confirming reproducibility, differential representation of individual sgRNAs in TI- vs. P-Tregs was significantly correlated in the two cohorts (*p* < 0.01, Figure 2D). Of the 17 candidate MR proteins, 8 presented significantly depleted sgRNAs in TI-Tregs vs. spleen P-Tregs, including—TRPS1, MAFB, BANP, FOSL2, EGR3, GLI1, NR3C1, and ZEB2 (Figure 2E), suggesting a causal role in Treg tumor infiltration and/or retention to the TME. Critically, 6 of 8 validated MR proteins—including TRPS1, MAFB, FOSL2, EGR3, GLI1, and ZEB2—were also significantly depleted in TI-Tregs relative to tumor CD4 Tconv, (Figure 2F), thus supporting their Treg-specific function. When restricted to validated genes, differential sgRNA representation was even more significantly correlated across mouse cohorts (p = 6.0×10^-4^, Figure S2B-C). Confirming the quality of the data, the sgRNAs of the positive control Foxp3, which is Treg but not CD4 Tconv essential, were differentially represented in the two populations (*p* = 8.68×10^-4^, Figure 2F).

The most statistically significant protein emerging from the study—in terms of TI-Tregs vs both P-Tregs and Tumor CD4 Tconv depletion (p=2.21×10^-175^ and p=1.72×10^-76^, respectively)—was TRPS1, a protein with unknown function in T-cells, including Tregs.

### Loss of TRPS1 in Hematopoietic Lineages Inhibits Tumor Growth

Based on these findings, we focused on TRPS1, the MR whose guide RNAs were most significantly depleted in TI-Tregs vs. both P-Tregs and Tumor CD4 Tconv. Specifically, two guide RNAs targeting the encoding gene, *Tprs1*, were transduced into Cas9-expressing LSKs that were then used to reconstitute the bone marrow of six lethally-irradiated chimeras. As negative controls, we reconstituted the bone marrow of a 5-mouse cohort with LSKs transduced with two non-targeting (*scramble*) guides. To assess the tumor-agnostic nature of TI-Treg infiltration MRs, we implanted these mice with a different syngeneic tumor model, MCA205, representing a well-studied, poorly immunogenic fibrosarcoma (Pfirschke et al., 2016). Confirming the MR’s functional relevance, we observed a significant survival advantage in *Trps1*-ko mice vs. controls (p = 0.047) (Figure 2G-I). In particular, we observed spontaneous, durable tumor rejection (> 60 days) in three of the six *Trps1*-ko animals (Figure 2I). We hypothesize rejection was not observed in all Trps1-ko animals due to incomplete knockout efficiency in our system that may have introduced variability in the readout (25-40% maximum sgRNA transduction efficiency; Figure S2A). However, taken together, these data suggest that TRPS1 activity is essential for Tregs to maintain their immunosuppressive potential in the TME.

### Systematic Identification of TI-Treg-specific MR-inverter Drugs

To identify mechanism-based drugs that could specifically inhibit Treg infiltration/retention to the TME by targeting the MR proteins identified by our study, we generated RNA-seq profiles of human-derived TI-Tregs at 24h following treatment with a library of clinically relevant compounds. To reduce study complexity and cost, we first assessed the effect of a library of 1,554 FDA-approved and investigational compounds on human-derived P-Treg viability at a single, relatively large concentration (5 μM). For this screen, human P-Tregs were flow sorted, expanded *ex vivo*, and drug treated in a 96-well plate format (Figure 3A). We then selected 195 compounds that inhibited P-Treg viability ≥ 60% (Figure 3B). To further reduce the number of candidate drugs, we then generated 10-point drug response curves to identify the 48h EC_20_ concentration of the 195 compounds, then selected a repertoire of 86 compounds with the lowest EC20 for efficient perturbational profile analysis in a 96-well format, when considering inclusion of vehicle controls (DMSO). As previously reported (Bush et al., 2017; Douglass et al., 2020), the 48h EC_20_ (*maximum sublethal*) concentration was selected to effectively assess the 24h drug mechanism of action, while reducing confounding effects arising from activation of cell stress or death pathways. Finally, the EC_20_ concentration of each compound was then used to perturb TI-Tregs flow-sorted from a chemo-naïve human clear cell carcinoma specimen and expanded *ex vivo,* into 96-well plates, followed by RNA-seq profiling using the fully automated PLATE-Seq technology (Bush et al., 2017; Douglass et al., 2020). Viability, as well as perturbational RNA-seq profiles of TI-Tregs were collected (Figure 3C-D). By using the differential protein activity signature in drug- vs. vehicle control-treated TI-Tregs, we identified the compounds capable of inducing the most statistically significant inactivation of TI-Treg-specific MR proteins (OncoTreat algorithm (Alvarez et al., 2018). From this analysis, 32 compounds were nominated as statistically significant inhibitors of the 17-MR protein signature (Figure S3B), seven of which preferentially depleted TI-Treg vs. P-Treg viability *in vitro* (Figure 3C-D), and which were also found to inhibit patient-by-patient TI-Treg vs P-Treg MRs across all tumor types and nearly all patients (Figure 3E). Of these, three (*i.e.*, gemcitabine, triapine, and floxuridine) were among the seven most significant TI-Treg MR activity inhibitors—as assessed from the individual TI- vs. P-Treg signatures of all 36 patients in the study (Figure S3A)—and among the top 6 inducing the most significant differential TI- vs. P-Treg viability reduction *in vitro* (Figure 3C).

**Figure 3:**
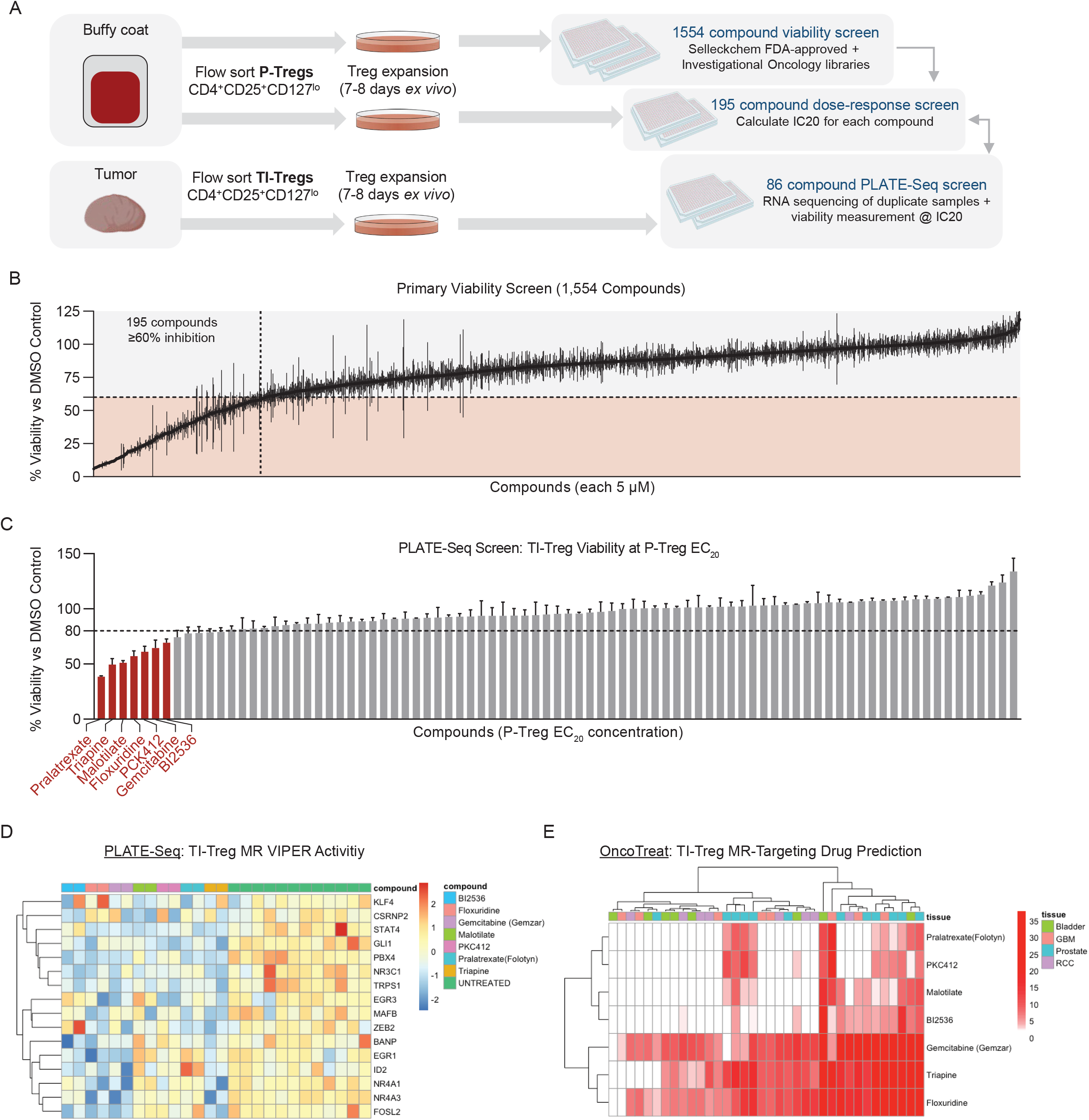
High-Throughput Drug Screening Platform Identifies Potential Drug Candidates With Tumor-Treg-Directed Toxicity: **(A)** Experimental design of High-Throughput Treg-Directed Drug Toxicity Screen. **(B)** Results from initial set of 1,554 FDA-approved and investigational Oncology compounds screened at single-dose for peripheral Treg growth inhibition, with 195 compounds showing >60% inhibition at 5uM. **(C)** Viability results of the PLATE-Seq screen, where human tumor Tregs were assessed for growth inhibition on sorted Tumor Tregs at peripheral-Treg EC20 dose, resulting in 7 drugs with higher toxicity in TI-Tregs relative to P-Tregs. **(D)** Heatmap of VIPER protein activity for Tumor vs Peripheral Treg MRs defined in 1E, 1F comparing transcriptional effect of drugs in (C) vs untreated control, with downregulation of nearly all identified Master Regulators by these drugs. **(E)** Patient-by-Patient Drug predictions according to inversion of patient Tumor Treg vs Peripheral Treg protein activity signature by drug-treatment protein activity signature. Each drug predicted to invert Tumor Treg signature with -log10(Bonferroni-Corrected p-value) < 0.01 in a particular patient is colored red. Patients are grouped by tumor type. Subset to drugs identified by tumor Treg growth screen in (C), with columns colored by tumor type and clustered by unsupervised hierarchical clustering.

Dose response curves of these three drugs revealed that only gemcitabine had a gradual effect on Treg viability reduction, as a function of its concentration, while the other two had sharp elbows that would challenge appropriate concentration selection for in vivo studies (Figure S4A-C). In addition, floxuridine had cytostatic rather than cytotoxic activity even at high concentration. Surprisingly, gemcitabine elicited differential TI-vs. P-Treg sensitivity *in vitro* at a concentration of only 10nM (Figure S3B), which is much lower than the clinically relevant concentration. As a result, we focused on this drug for *in vivo* validation purposes.

### Low-Dose Gemcitabine inhibits tumor viability preferentially in immunocompetent mice

To validate the preferential TI-Treg targeting of gemcitabine *in vivo*, we implanted C57BL/6J mice subcutaneously with the MC38 syngeneic tumor model, and initiated therapy 12 days later, a “late stage” of growth when MC38 tumors are reported to be broadly resistant to anti-PD-1 immunotherapy (Taylor et al., 2019). Gemcitabine was administered intra-peritoneally (IP) on days 12, 15, and 18, at 12 mg/kg, representing a low dose of 1/10^th^ of the lowest conventional clinical-equivalent dose in mice (120 mg/kg) (Ager et al., 2021; Beatty et al., 2011). Additional mouse cohorts received gemcitabine in combination with anti-PD-1 administered IP on days 12, 15, and 18 (Figure 4A). As expected, late stage MC38 tumors failed to respond to anti-PD-1. However, single agent low-dose gemcitabine temporarily controlled MC38 progression, conferring a significant reduction in growth kinetics (p=0.003) and prolongation of survival (p=0.006) relative to vehicle treated mice (Figure 4A-D). In combination, low-dose gemcitabine sensitized late stage MC38 tumors to anti-PD-1, achieving complete responses in 50% of animals, translating to a significant survival advantage compared to gemcitabine alone (p=0.009) (Figure 4A-D).

**Figure 4:**
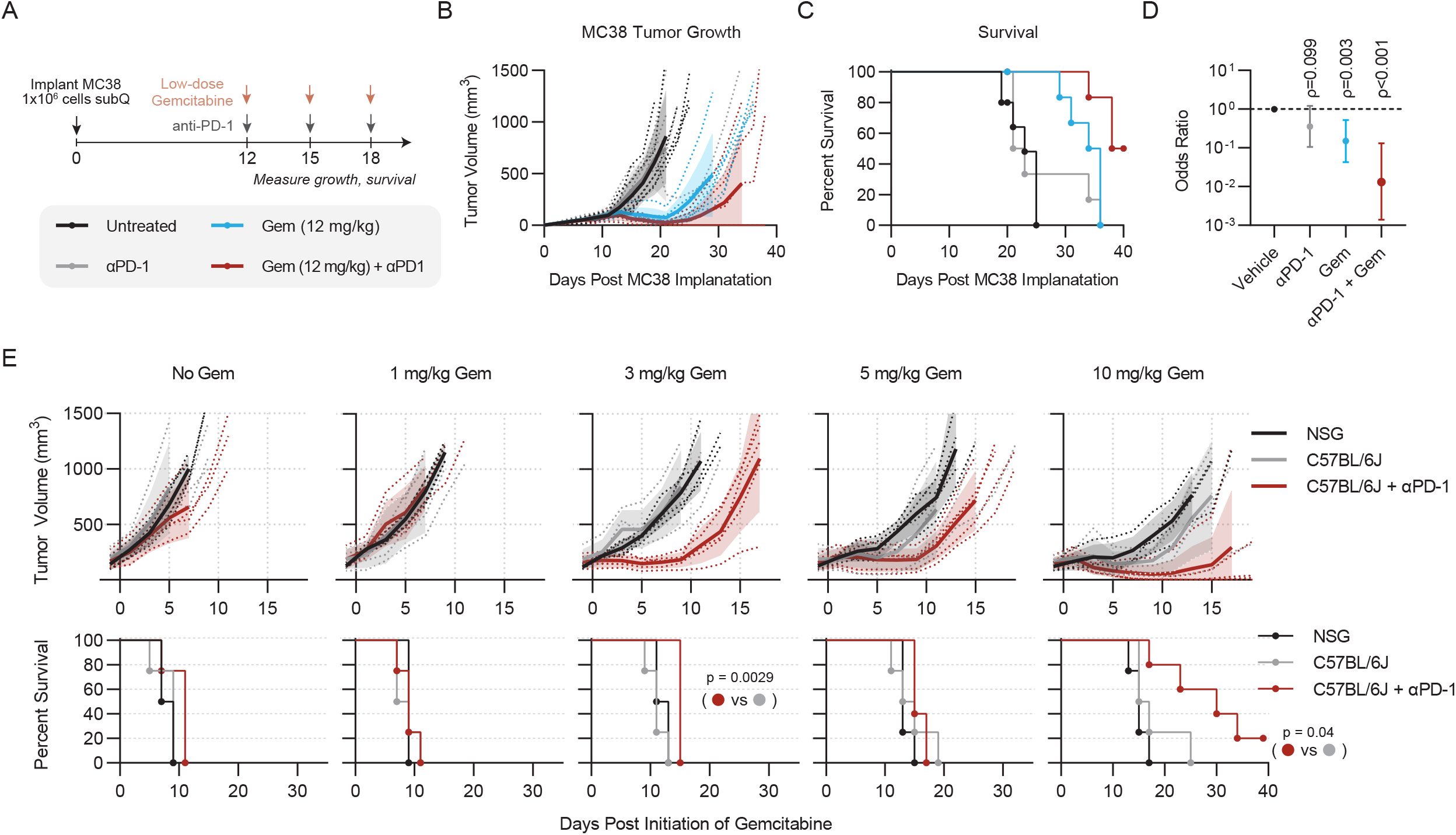
Low-Dose Gemcitabine is Immunogenic and Potentiates anti-PD-1 Therapy: **(A)** Schematic of *in vivo* validation studies.. Experiment consists of 6 mice per group. **(B)** Tumor growth curves for each treatment group, **(C)** Kaplan-Meier survival curves, and **(D)** forest-plot showing the result of multiple cox regression assessing treatment effect on time-to-death for each of the treatments described in (A). Hazard ratios are shown with 95% confidence interval and p-value, such that anti-PD-1+Gemcitabine most improves survival, followed by Gemcitabine monotherapy. Results are representative of two independent experiments. **(E)** Tumor growth and Kaplan Meier survival curves of NSG mice, C57BL/6J mice, and C57BL/6J mice exposed to anti-PD-1 therapy receiving the indicated dose of gemcitabine between 1-10 mg/kg. Statistical significance for survival was calculated by Mantel-cox log rank test.

To assess whether low-dose gemcitabine effects were immune-mediated, we performed parallel dose titrations in immune-competent C57BL/6J mice and severely immune-deficient NSG (NOD.Cg-Prkdc^scid^ Il2rg^tm1Wjl^/SzJ) mice lacking both innate and adaptive immunity. At the clinically-typical dose (120 mg/kg) (Ager et al., 2021; Beatty et al., 2011) gemcitabine inhibited tumor growth in both C57BL/6J and NSG mice relative to vehicle control (p < 0.001, by Cox regression analysis) with no significant difference between the two strains (*p* = 0.19, Figure S4E-F). We found efficacy was lost in both strains between the range of 12 mg/kg and 1.2 mg/kg, with a trending but non-significant advantage in C57BL/6 mice at 1.2 mg/kg (p=0.09 Figure S4F), suggesting immune-dependent effects of gemcitabine may be observed within this range.

To test whether immune-dependent activity could be observed in this range of concentrations, we dosed cohorts of mice with 1-10mg/kg gemcitabine, with an additional cohort of C57BL/6J mice receiving anti-PD-1 in combination. We found that doses as low as 3 mg/kg, which lack any activity in NSG mice (p=0.84), reveal sensitivity to anti-PD-1 via tumor growth kinetic reduction (p=0.01) and enhanced survival (*p* = 0.0029) in the combination group. At 10 mg/kg, we observe a trending survival advantage in the C57BL/6J vs. NSG strains (p=0.253) however in immune-competent mice this dose is sufficient to augment anti-PD-1 therapy to achieve curative responses and a significant enhancement of survival relative to gemcitabine or anti-PD-1 alone (p=0.048, p=0.005, respectively). Taken together, these data show that low-dose gemcitabine, while ineffective in the absence of host immunity, sensitizes anti-PD-1 resistant MC38 tumors to immune checkpoint blockade therapy.

Finally, to test the hypothesis that low-dose gemcitabine achieves tumor growth inhibition by modulating Treg recruitment/retention to the TME, we generated scRNA-seq from MC38 tumor- and spleen-derived Tregs, at 24 hours after treatment with a single 12 mg/kg dose of either gemcitabine or vehicle control (Figure 5A). For this study, we implanted *FoxP3^Yfp-Cre^* mice with MC38 tumor cells to facilitate specific flow-sorting of CD4^+^ FoxP3^+^ Tregs from tumor and spleen using YFP as a FoxP3 expression marker. Using a 5-mouse cohort per group, we obtained high quality profiles from ∼10,000 spleen-derived and ∼1,000 tumor-derived Tregs from each group (Figure 5B, S5A). Protein activity-based cluster analysis stratified the cells into five clusters (TRC_1_ – TRC_5_) (Fig 5C, S5B), with cluster TRC_3_ highly enriched for human TI-Treg MRs (Figure 5D-E). In vehicle-treated control animals, the TRC_3_ cluster comprised 7.8% of splenic Tregs vs. 30.1% of TI-Tregs (*p* = 1.8×10^-84^). Gemcitabine treatment reduced this TRC_3_ occupancy by ∼50%, to 14.9% of the TI-Treg cells, while inducing virtually no change in the spleen population (Figure 5F-G). Furthermore, treatment resulted in a proportional increase in TRC_1_ occupancy, which exhibits signs of interferon exposure (high IFI16 activity). These data suggest low-dose Gem has antagonistic effects on TI-Tregs presenting activation of TI-Treg MRs *in vivo*.

**Figure 5:**
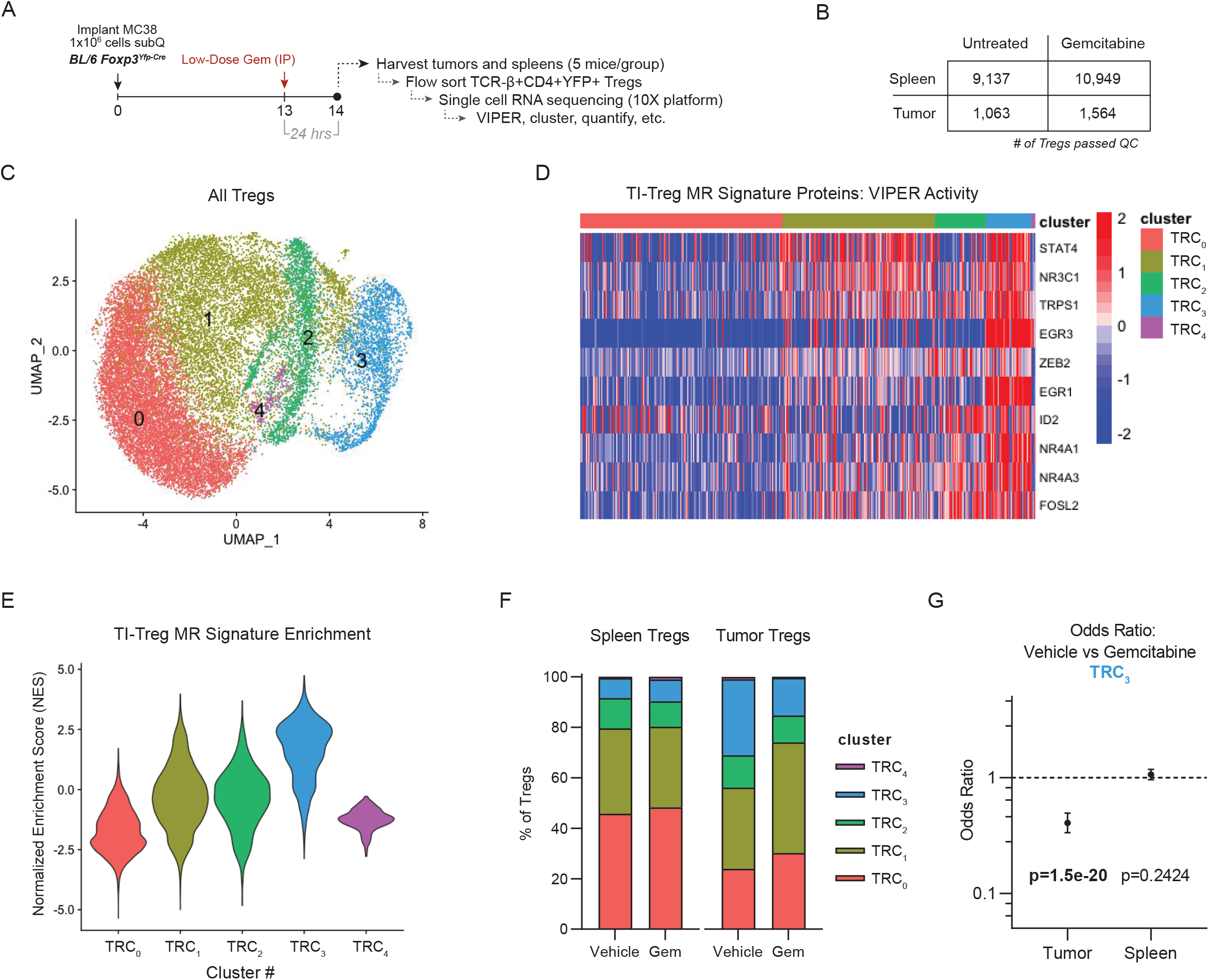
Single-Cell RNA-Sequencing Suggests Low-Dose Gemcitabine Depletes TI-Tregs. Exhibiting high TI-Treg Master-Regulator Activity: **(A)** Schematic of experimental workflow. **(B)** Numbers of Tregs successfully sequenced in each treatment condition and utilized in downstream analysis. **(C)** UMAP plot and unsupervised clustering by VIPER-inferred protein activity of Tregs from Untreated and Gemcitabine-treated Tumor and Spleen. **(D)** Heatmap of cell-by-cell protein activity for each Tumor-Treg MR identified by single-cell RNASeq, grouped by cluster. **(E)** Distribution of TI-Treg MR signature normalized enrichment score by Gene Set Enrichment Analysis (GSEA), grouped by cluster, such that cluster TRC_3_ is most enriched for the TI-Treg MR signature. **(F)** Barplot of cluster frequencies in each sample, such that cluster 3 has a baseline frequency of 7.8% in spleen of vehicle-control sample and 30.1% in tumor (p = 1.78e-84), with frequency of only 14.9% in tumor of gemcitabine-treated sample (p=1.51e-20). **(G)** Cox proportional hazard ratios of cluster 3 frequencies in vehicle vs gemcitabine treated mice in tumor (OR = 0.407 [95% CI: 0.334-0.494]) and spleen (p=0.242, OR = 1.063 [95% CI: 0.958-1.17]).

## Discussion

Treg immunosuppression in the TME is a major barrier to antitumor immunity and undermines efficacy of checkpoint blockade immunotherapy, which remains effective only in a minority of cancer patients (Chao and Savage, 2018; Zappasodi et al., 2018). To address the critical need for more effective agents to counteract human TI-Treg number or function, we harnessed new tools to identify and validate previously unappreciated regulators of TI-Tregs. Applying VIPER protein activity inference on a novel dataset of TI-Tregs, P-Tregs, and additional CD4 and CD8 non-Treg controls across 36 patients, we defined a set of 17 TI-Treg MRs (EGR1, NR3C1, PBX4, MAFB, ID2, STAT4, NR4A3, NR4A1, TRPS1, EGR3, BANP, ZEB2, KLF4, GLI1, CSRNP2, KDM2B, FOSL2), which we then functionally validated in a pooled *in vivo* CRISPR screen. First among the validated targets was TRPS1, a transcription factor not previously studied in the context of Treg biology, which was also found to be the most strongly suppressive of TI-Tregs relative to tumor infiltration by CD4 Tconv cells. In parallel, we conducted a systematic *ex vivo* drug screen and found gemcitabine possesses preferential cytotoxic capacity against TI-Tregs and inhibits transcriptional activity of TI-Treg MRs, including TRPS1, across multiple tumor types. With *in vivo* validation studies we found sub-clinical doses of gemcitabine that lack full activity in immune deficient animals are sufficient to potentiate checkpoint blockade immunotherapy control of established, anti-PD-1 resistant MC38 tumors. These findings have implications for both basic understanding of TI-Treg biology as well as clinical use of available chemotherapeutics for the purpose of modulating TI-Treg activity.

Our findings showing immune-modulating properties for gemcitabine are broadly consistent with prior reports. Early studies showed that clinically equivalent doses of gemcitabine systemically decrease MDSC and B cell numbers without substantial effects on T cells, and in fact promote T cell trafficking into tumors (Daikeler et al., 1997; Nowak et al., 2002; Suzuki et al., 2005). In multiple pre-clinical models, tumor growth in T cell-deficient Nude mice or specific CD8 T cell depletion rendered gemcitabine less effective, suggesting that gemcitabine exhibits T cell-dependent immunogenic activity in addition to direct tumoricidal killing (Suzuki et al., 2007). Informed by prior investigation into immunogenic effects of low dose or metronomic dosing of other chemotherapeutic agents such as cyclophosphamide (Ghiringhelli et al., 2007; Lutsiak et al., 2005; Wada et al., 2009) or oxaliplatin (Shalapour et al., 2015), more recent studies showed that sub-clinical “low” doses of gemcitabine are immunomodulatory in various ways, with effects on NK cell function (Zhang et al., 2020), myeloid polarization (Deshmukh et al., 2018; Di Caro et al., 2016) and Tregs (Homma et al., 2014; Rettig et al., 2011; Shevchenko et al., 2013; Skavatsou et al., 2021; Tongu et al., 2013).

While in agreement with the observation that gemcitabine antagonizes Tregs in mouse and man, our findings clarify that, at low doses, gemcitabine is preferentially toxic to TI-Tregs as compared to P-Tregs, this finding was not clear in prior studies. That finding also extends our understanding of gemcitabine’s mechanism of action to include inhibition of key TI-Treg master regulators, particularly TRPS1. Future studies are required to more fully understand how gemcitabine selectively modulates TI-Treg MRs. Furthermore, our gemcitabine titration studies in immunocompetent C57BL/6 versus severely immunodeficient NSG mice defined a more narrow “low dose” range at which gemcitabine is primarily immunomodulatory, building upon previous studies in Nude mice that were confounded by the presence of functional NK and myeloid cells, as these are also known to be modulated by gemcitabine (Suzuki et al., 2007; Zhang et al., 2020). We found the range between 3-10 mg/kg of gemcitabine dosed Q3D to be immunogenic, which represents 2.5-8.3% of the standard murine maximum tolerated dose of 120 mg/kg, and roughly translates to a human equivalent dose of 9-30 mg/m^2^ (Nair and Jacob, 2016), as compared to the standard clinical dose of 1,000 mg/m^2^. Although we fully acknowledge the challenges of translating dosing strategies between species, our studies support the development of dose-finding studies of gemcitabine in combination with immune modulating agents such as anti-PD-1, particularly in settings where the benefit of anti-PD-1 monotherapy is sub-optimal.

A major finding of our study was the discovery and validation of TRPS1 as a putative master regulator of TI-Tregs, such that TRPS1-ko inhibits tumor Treg infiltration without depleting peripheral Tregs, preferentially inhibits TI-Tregs relative to tumor CD4 Tconv, and inhibits overall tumor growth, with a 50% cure rate in MCA205 tumor model. TRPS1 is a transcription factor classically linked to skeletal development, as subjects with germline alterations in the *Trps1* gene suffer from autosomal dominant trichorhinophalangeal syndromes with characteristic craniofacial abnormalities (Lüdecke et al., 2001; Momeni et al., 2000). More recently, TRPS1 has been implicated in tumorigenesis in breast cancer (Ai et al., 2021; Cornelissen et al., 2020) and osteosarcoma (Li et al., 2015) potentially through promotion of dysregulated cell replication resulting in accumulation of genomic aberrations (Yang et al., 2021). Functionally, TRPS1 is thought to function uniquely as a transcriptional repressor via its GATA domain (Malik et al., 2002), although notably TRPS1 contains two Ikaros-like domains whose specific functions are poorly characterized. Other Ikaros family proteins including Helios and Aiolos are expressed in hematopoietic tissues with important functions in Treg differentiation and function (Getnet et al., 2010; Zabransky et al., 2012), thus it is tempting to speculate that TRPS1 governs TI-Treg activities via its Ikaros domain. At this point, the specific functions of TRPS1 in Tregs remain to be described and additional future work is warranted to understand mechanisms of TRPS1 regulation in Tregs both within and outside of the tumor microenvironment. Additionally, our results support the design of specific inhibitors of TRPS1 activity, given its putative role in Tregs, high activity in TI-Tregs across multiple cancer types, and effect on tumor growth, as well as its known cell-intrinsic pro-tumorigenic role in multiple cancer types (Ai et al., 2021; Cornelissen et al., 2020; Li et al., 2015).

Together, the approach here – combining CRISPR validation of putative regulatory proteins in an *in vivo* functional genomics system with *ex vivo* drug screening and transcriptional profiling of treatment response represents a novel approach to the discovery of Treg-directed immunotherapy targets. Of note, this platform could in theory be extended to other immunosuppressive hematopoietic cell types, opening up additional possibilities for target identification and validation across the field of immuno-oncology. Furthermore, our PLATE-Seq screening method can be feasibly extended to a significantly larger number of compounds, allowing for discovery of additional compounds with preferential activity against TI-Tregs. The utility of this approach is exemplified by our discovery low-dose gemcitabine as an TI-Treg modulating agent, which we have newly found to function at least in part through antagonism of TI-Treg master regulator programs, including inhibition of a novel TI-Treg regulator TRPS1. While the development of TRPS1-directed therapeutics will require additional effort, our findings on low-dose gemcitabine are readily translatable to human studies aimed at improving the clinical activity of anti-PD(L)-1 agents in the clinic.

## Methods

### Clinical Sample Collection, Sorting, and RNA-Sequencing

Tissue was collected from treatment-naïve resected tumors across patients with four tumor types, including 8 patients with glioblastoma multiforme, 8 patients with clear cell renal carcinoma, 8 patients with bladder cancer, and 12 patients with prostate cancer. For each patient, 50ml of peripheral blood was drawn at the same time that tumor was resected. Tumors were dissociated with the GentleMACS OctoDissociator following manufacturer’s instruction, and subsequently Tregs and CD8 T-cells were flow-sorted from tumor along with Tregs, naïve CD4nonTregs, and naïve CD8 T cells from peripheral blood. An aliquot of naïve CD8 and CD4nonTreg were stimulated ex vivo with IL2 and anti-CD3/anti-CD28 beads for 24 hours to induce T-cell activation. Flow-sorted and ex-vivo-stimulated populations were processed to prepare cDNA libraries following Illumina user guide and were sequenced on Illumina NovaSeq 6000 Sequencing System.

### Gene Expression and VIPER Analysis

Gene Expression was combined across all samples and scaled to log10(Transcripts Per Million + 1). Gene Expression was subsequently scaled across rows by z-score transformation and used as input for Principal Component Analysis (Figure 1A) and differential gene expression.

Log10(TPM+1) matrix was separately used to infer gene regulatory network structure by the ARACNe algorithm. ARACNe was run with 100 bootstrap iterations using 1785 transcription factors (genes annotated in gene ontology molecular function database as GO:0003700, “transcription factor activity”, or as GO:0003677, “DNA binding” and GO:0030528, “transcription regulator activity”, or as GO:0003677 and GO:0045449, “regulation of transcription”), 668 transcriptional cofactors (a manually curated list, not overlapping with the transcription factor list, built upon genes annotated as GO:0003712, “transcription cofactor activity”, or GO:0030528 or GO:0045449), 3455 signaling pathway related genes (annotated in GO biological process database as GO:0007165, “signal transduction” and in GO cellular component database as GO:0005622, “intracellular” or GO:0005886, “plasma membrane”), and 3620 surface markers (annotated as GO:0005886 or as GO:0009986, “cell surface”). ARACNe is only run on these gene sets so as to limit protein activity inference to proteins with biologically meaningful downstream regulatory targets, and we do not apply ARACNe to infer regulatory networks for proteins with no known signaling or transcriptional activity for which protein activity may be difficult to biologically interpret. Parameters were set to zero DPI (Data Processing Inequality) tolerance and MI (Mutual Information) p-value threshold of 10^-8^, computed by permuting the original dataset as a null model.

Using the ARACNe gene regulatory network structure, VIPER protein activity inference was performed on gene expression signature. First directly on z-score-scaled gene expression signature for all T-cell subtypes, used for Principal Component Analysis and clustering (Figure 1B). Then separately scaling Tumor and Peripheral Tregs against naïve CD4nonTregs by viperSignature command in Rstudio for comparison of Tumor Treg vs Peripheral Treg (Figure 1C, 1E), and scaling all Tregs and CD4nonTregs against naïve CD8nonTregs by viperSignature for comparison of Tumor Treg vs all Treg and CD4nonTreg controls (Figure 1D, 1F).

### Random Forest Feature Selection

The full dataset was randomly split into 75% training data and 25% testing data. On training data, a Random Forest Model was built with VIPER-inferred protein activity to classify Tumor Treg vs Peripheral Treg (Figure 1C) or Tumor Treg vs all Controls (Figure 1D), taking the list of all differentially active proteins (t-test p-value < 0.01) as an initial feature set. Features were ranked by mean decrease in model accuracy and included one-by-one to construct random forest models with feature selection. Predictive power was assessed by Area-Under-ROC-Curve (AUC) in the held-out testing data, and a null model of AUC was constructed from random sampling of the same number of genes (from the set of genes with differential activity p-value =1.0) 1000 times. For each comparison, the maximum number of discriminative genes was selected for which AUC vs null model remained statistically significant (Figure 1C, 1D). These genes are shown in Figure 1E and 1F and aggregated into a combined list of 17 putative Tumor Treg vs Peripheral Treg Master Regulators with Activity specifically upregulated in Tumor Tregs.

### CRISPR Validation in Chimeric Immune Editing Model

Confirmatory evidence that the predicted proteins regulate tumor Treg infiltration was generated in murine models in a pooled CRISPR screen (Figure 2); by comparing the differential representation of gene-knockout Tregs in tumor versus non-tumor tissue (spleen, as a control), for each candidate Master Regulator gene. For these studies, Hematopoietic Stem Cells (HSCs) were extracted from Cas9+ mice and transduced with sgRNA library targeting 34 genes with 3 guides/gene. The transduced stem cells were reimplanted into irradiated recipient mice, allowing reconstitution of the entire immune system, including Tregs, with a unique pool of CRISPR knockout genes in place. Subsequent implantation of a subcutaneous MC38 murine colon adenocarcinoma tumor model allowed direct observation of differential infiltration of tumors by Tregs receiving selected CRISPR guides, in a single, high-throughput experimental screen. Critically, the experiment would not have been possible on a genome-wide level without initial narrowing of candidate master regulators by VIPER protein activity analysis, due to fundamental limitations in achieving a sufficient number of tumor-infiltrating Tregs harboring guide DNAs for the full set of human genes. This is because we typically find fewer than 10,000 tumor-infiltrating Tregs in MC38 tumor model.

We designed the gDNA library with three guides per gene targeting the 17 predicted Tumor Treg MRs and 13 randomly sampled negative control genes (genes with p=1.0 comparing Tumor Treg to Peripheral Treg). We also included Treg context-specific positive controls such as Foxp3 and Cd4 and core-essential genes Cdk1 and Plk1 (these were not detected in any cells post-transduction, indicating successful gene-editing). For guide design, we used the Broad Institute Genetic perturbation platform (GPP) sgRNA designer-tool. Sorted Cas9+ hematopoietic stem cells were successfully transduced and implanted into irradiated recipient mice, A cohort of six replicate mice (cohort 1) and three replicate mice (cohort 2) were separately implanted and harvested, and Vex+ gDNA-bearing Tregs and CD4nonTregs were flow-sorted from Tumor and spleen, separately.

Pelleted Tregs/CD4s were first pooled together, with entire tumor samples pooled and spleen samples pooled in proportion to ratio of sorted cell counts from Tumor. gDNA was extracted first by adding 400ul of RIPA buffer (with added RNAseA) on top of the pelleted Tregs/CD4s, followed by 1h incubation at 65C. This was followed by Phenol/Chloroform/Isoamyl alcohol-extractions and Isopropanol-precipitations. Extracted gDNA was divided into 8 replicates with equal volumes for cohort 1 and four replicates with equal volumes for cohort 2, each then amplified by 2-step PCR and then sequenced. Correlation between replicates was assessed (Figure 2D).

After sorting and gDNA sequencing, differential frequency of guides in Tumor Treg vs Peripheral Treg and Tumor Treg vs Tumor CD4nonTreg are assessed by DESeq with Bonferroni correction on the p-values, separately (Figure S2B), and then p-values were integrated by Stouffer’s Method (Figure 2E-F).

### CRISPRko library design

For CRISPRko screening we designed the target gene list to include 34 genes (3 sgRNAs / gene)—including 17 MRs and 13 negative control genes (genes whose loss is not predicted to differentially affect Tumor Tregs compared to Peripheral Tregs i.e. p=1.0 comparing Tumor Treg to Peripheral Treg), and 4 positive controls (2 genes whose loss is known to be toxic to Tregs (FOXP3 and CD4) and 2 core-essential genes (PLK1 and CDK1)). Positive control sgRNAs were not detected in any cells post-transduction, indicating successful gene-editing. For guide design, we used the Broad Institute Genetic perturbation platform (GPP) sgRNA designer-tool. The pooled guide-library was ordered from Twist-bioscience. The guide sequences are found in (Supplementary table S1).

### CRISPRko oligo synthesis and library cloning

Oligo libraries (102 oligos) were ordered from Twist-biosciences in following format (200mers):

ACACGTCATATAGATGCCGTCCTAGCGAGCGTGGAGTGAGCCATTGTGAGCGCTCACAATTATATATCTTGTGGAAAGGACGAAACACCGNNNNNNNNNNNNNNNNNNNNGTTTTAGAGCTAGAAATAGCAAGTTAAAATAAGGCTAGTCCGTTATCATCGGCAGCAACCAGATGGGCACAGGAAAGATACTTAACGCTT

From the initial oligo pool, this TREG sub-library was amplified first with KAPA polymerase (KK2502) with the following PCR primers and settings:

**TREG_1F:** AGCGTGGAGTGAGCC,

**TREG_1R:** TCTGGTTGCTGCCGA

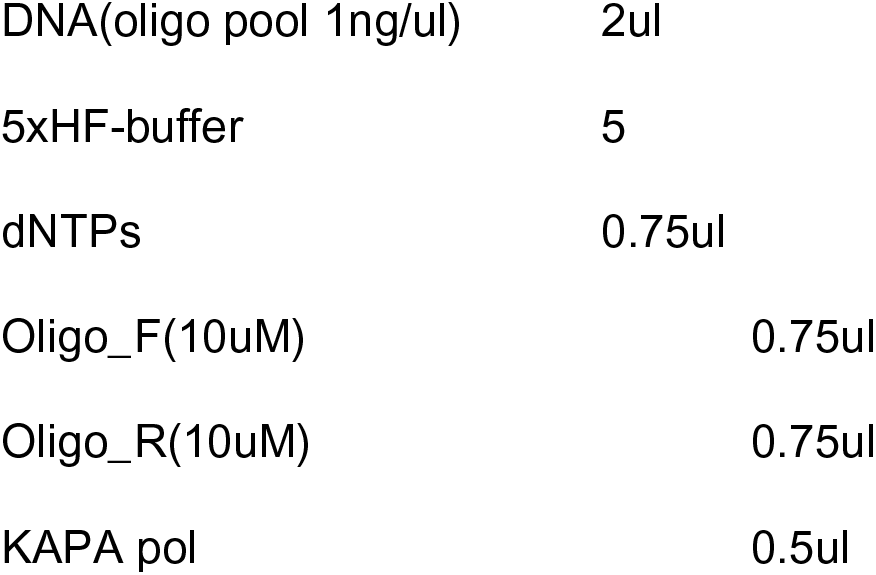
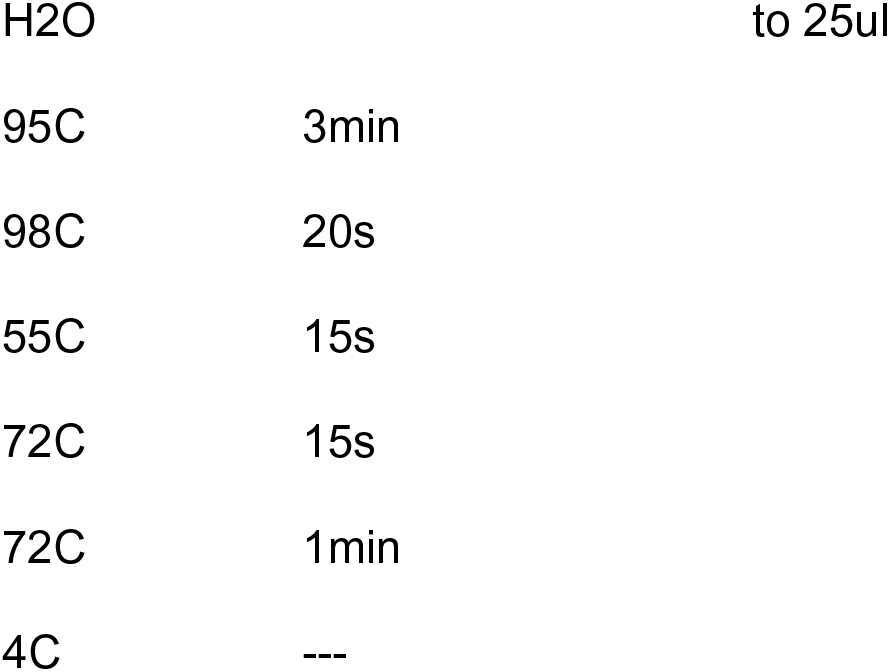

|  |  |
| --- | --- |
| DNA(oligo pool 1ng/ul) | 2ul |
| 5xHF-buffer | 5 |
| dNTPs | 0.75ul |
| Oligo_F(10uM) | 0.75ul |
| Oligo_R(10uM) | 0.75ul |
| KAPA pol | 0.5ul |

|  |  |  |
| --- | --- | --- |
| H2O |  | to 25ul |
| 95C | 3min |  |
| 98C | 20s |  |
| 55C | 15s |  |
| 72C | 15s |  |
| 72C | 1min |  |
| 4C | --- |  |

The PCR product from PCR1 was gel purified with GeneJet gel purification-kit. The final 2nd PCR prior to the Gibson cloning-step was done with the following primers and settings:

**TREG_2F:** AGCGCTCACAATTATATATCTTGTGGAAAGGACGAAACACCG

**TREG_2R:** CGGACTAGCCTTATTTTAACTTGCTATTTCTAGCTCTAAAAC

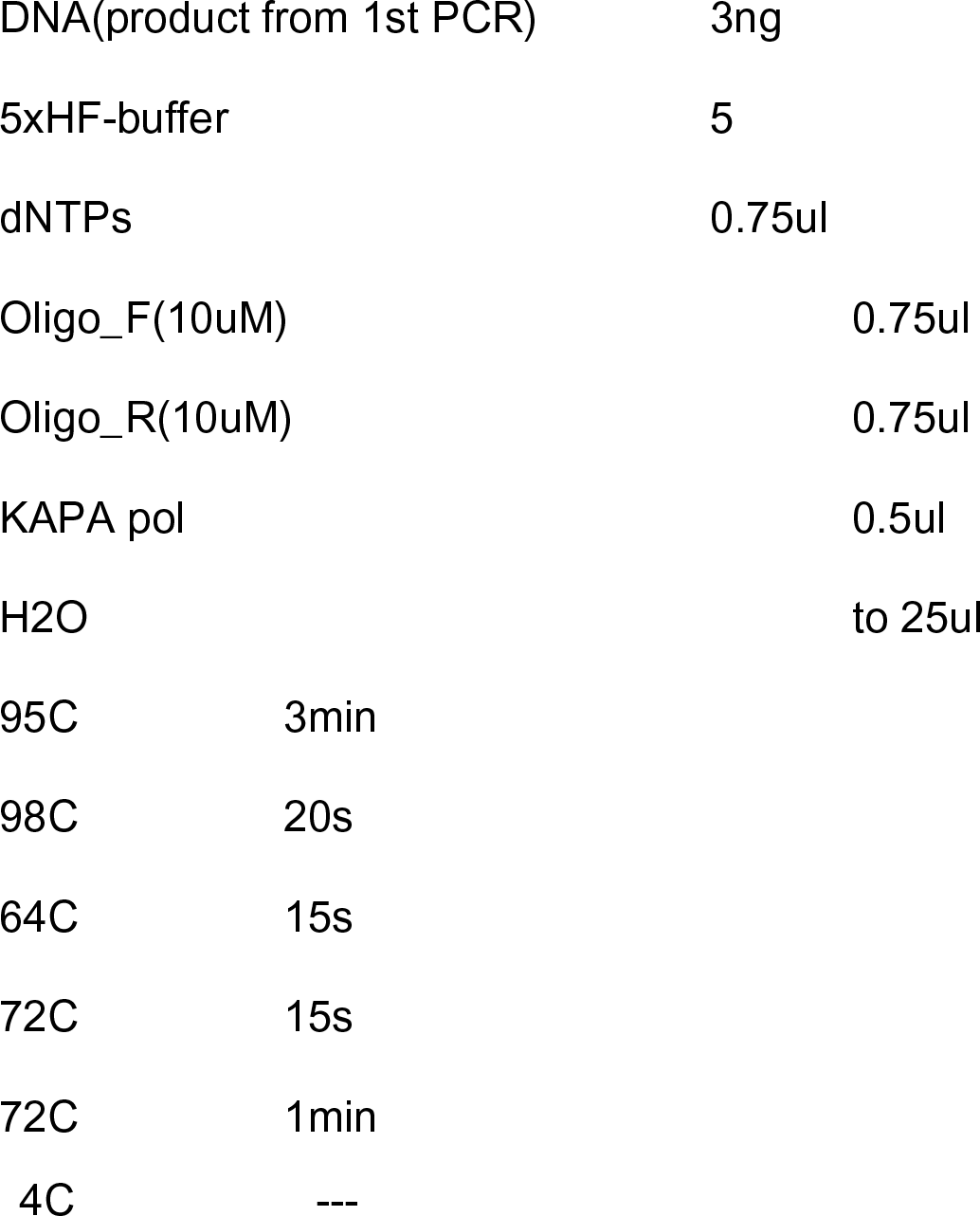

Both of these amplifications were done with qPCR and the program was stopped before the amplification started to plateau. After PCR the insert was gel purified (GeneJet) and Gibson cloned into BsmBI-digested pXPR_053 (Addgene# 113591). Gibson cloned insert and vector was column purified (GeneJet) and large-scale electroporated into Lucigen Enduro competent cells. The bacterial colonies were scraped from 24,5cm x 24,5cm agar plates, so that the estimated library complexity was approximately 1000 colonies / sgRNA.

### Lentiviral packaging of the sgRNA library

13 million 293T cells were seeded for each 15cm dish the night before transfection. The following morning, viral transfections were conducted with the following components:

- 22.1ug sgRNA containing pXPR_053 (Addgene 113591).
- 16.6ug PsPAX2 (Addgene 12260)
- 5.5ug PMD2G (Addgene 8454).
- 1660ul of sterile H2O.

After mixing the plasmids and H2O, 110,6ul of Fugene HD (Promega) was added to the mix. The transfection mixture was vortexed, then incubated for 10 minutes before adding dropwise to 293T cells. The transfection mixture was removed the following day and virus was collected at 48h and 72h after initial transfections. To remove cellular debris, the virus-containing supernatant was centrifuged 500 x g for 5min and filtered with 0.45um PES filters (Millipore), followed by ultracentrifugation (25,000rpm for 2h), dissolving the viral pellet into PBS, aliquoting the virus and storing the aliquots at -80C. Viral titer was measured with 293T cells by using violet-excited GFP in the pXPR_053-plasmid.

### Cell culture and sgRNA transductions into hematopoietic LSK cells

HEK293T cells: HEK293T cells used in this study were obtained from the American Type Culture Collection (ATCC) and cultured at 37 °C in a humidified incubator (5% CO2) with the following media: DMEM + 10% FBS, 1% L-Glutamine and 1% Penicillin/Streptomycin. Cell line was tested for mycoplasma status before viral production. LSKs: After sorting the LSKs from donor mice, cells were sorted into 96-well plate (100k LSKs/well) and incubated overnight in SFEM media supplemented with 100 ng/mL of the following cytokines: SCF, TPO, Flt3-Ligand, and IL-7. Pen/Strep was used in all in vitro cultures. The following day, LSK cells were transferred into Retronectin-coated 24-well plate and sgRNA-containing Lentiviruses were added to the wells with MOI 30 (based on viral tittering in 293T cells). The final volume was adjusted to 400ul / well by adding cytokine supplemented SFEM stem cell media. The cells were centrifuged at 650 x g for 1.5 hours at 37°C with an acceleration of 2 and a brake of 1. After centrifugation, the plate was placed into 37C incubator for 1h, before adding 500 microliters of prewarmed stem cell media on top of the LSKs and overnight incubation. Transduced LSKs were implanted into donor mice irradiated with two doses of 600rads, spaced four hours apart, by intravenous tail vein injection immediately following the second irradiation.

### Genomic DNA extraction

Since the number of Vex+ tumor Tregs was very low in any individual mouse and because the mice all share the same genetic background, we decided to pool all tumor Tregs and tumor CD4s together across mice before the gDNA extraction step in order to reliably purify gDNA with sufficient yield. After the gDNA extractions, the extracted gDNA was split evenly into 8 (for cohort 1) or 4 (for cohort 2) separate technical replicates and library prep PCRs and NGS were done individually to all these technical replicates. In other words, genomic DNA was extracted by pooling all the FACS sorted Vex+ tumor Tregs (or tumor CD4 cells) from all the mice within each cohort and lysing the cells with 400ul of RIPA-buffer + RNAseA, followed by 1h incubation in 65C. After this, 400ul of Phenol/Chloroform/Isoamyl alcohol was added, followed by 6 min centrifugation at room temperature. Finally, the gDNA was recovered by Isopropanol precipitation. For spleen Tregs and spleen CD4 all the gDNA extractions were done individually for each mouse-sample (not pooled together at the lysis-stage as with tumor Tregs and tumor CD4s), since the number of extracted Vex+ cells was much higher than with tumor Tregs / CD4s. Otherwise, the protocol was identical compared to gDNA extractions from tumor Tregs and CD4s.

### Preparation of NGS libraries from the extracted gDNA

NGS libraries were prepared from extracted gDNAs following a 2-step PCR protocol with 2 x KAPA Mastermix (KK2612, KAPA Biosystems). For spleen Tregs and CD4s, individually purified gDNAs were pooled before the NGS library prep PCRs. This was done by pooling Spleen Tregs and CD4s in the same ratio as Tumor Tregs and CD4s previously pooled for gDNA extraction as measured by Vex+ FACS cell count. Before the 1st PCR, all pooled Treg and CD4 samples were split into 8 or 4 (first and second cohort) technical replicates, which were amplified separately and with different sample indexes. Correlation between replicates by gDNA frequency was assessed in each cohort and for each set of replicates following library sequencing (Figure 2D). Both 1st and the 2nd PCRs were stopped before amplification started to saturate in order to avoid biases in the library coverage. The following primers and PCR programs were used for the NGS library preps:

**TREG_NGS_1F:** GGACTATCATATGCTTACCGTAACTTGAAAGTAATTGT

**TREG_NGS_1R:** GAAGATCCGGGTGACGCTGCGAACGGACGT

**1st PCR:**

|  |  |
| --- | --- |
| gDNA | 12.5 - 25% of pooled material (depending on the cohort) |
| 2 x KAPA mastermix | 12.5ul |
| Oligo_F(10uM) | 1ul |
| Oligo_R(10uM) | 1ul |
| H2O | to 25ul |
| 95C | 3min |
| 98C | 20s |
| 60C | 15s |
| 72C | 20s |
| 72C | 1min |

**TREG_NGS_2F:**

AATGATACGGCGACCACCGAGATCTACACTCTTTCCCTACACGACGCTCTTCCGATCT(0-8nt stagger)TTGTGGAAAGGACGAAACACCG

**TREG_NGS_2R:** CAAGCAGAAGACGGCATACGAGATNNNNNNNNGTGACTGGAGTTCAGACGTGTGCTCTTCCGATCTTCTACTATTCTTTCCCCTGCACTGT

|  |  |
| --- | --- |
| 1:50 diluted DNA template from PCR 1 | 8ul |
| 2 x KAPA mastermix | 12.5ul |
| Oligo_F(10uM) | 1ul |
| Oligo_R(10uM) | 1ul |
| H2O | to 25ul |
| 95C | 3min |
| 98C | 30s |
| 52.5C | 15s |
| 72C | 20s |
| 72C | 1min |

After the 2nd PCR, samples were gel-purified (GenJet), pooled and sequenced with Illumina. Sequencing reads were aligned to a reference of sgRNA template sequences by kallisto to determine a counts matrix of reads per guide for each sample. Differential frequency of guides targeting the same gene in Tumor Treg vs Peripheral Treg and Tumor Treg vs Tumor CD4nonTreg was assessed by DESeq with Bonferroni correction on the p-values, separately (Figure S2B-C), and then p-values across cohorts were integrated by Stouffer’s Method (Figure 2E-F).

### High-Throughput Treg-Directed Drug Screening

From an initial library of 1,554 FDA-approved or investigational oncology compounds (SelleckChem), single-dose viability screening was performed in vitro on human Tregs sorted from Buffy Coat peripheral blood mononuclear cells (PBMCs). 195 compounds were identified which reduced peripheral Treg growth by at least 60% relative to DMSO control at 5uM. For these, dose-response titrations were performed to identify the IC20 dose at which peripheral Treg growth is inhibited by 20%, either by direct toxicity to Tregs or inhibition of Treg cell division. Subsequently, Tumor-Infiltrating Tregs were sorted from a large clear cell renal carcinoma tumor and plated with Treg-expansion beads in culture for 5 days, resulting in 5-million Tumor-Infiltrating Tregs. These were suspended at 160,000cells/mL and divided among 2 replicate plates for downstream RNA-Sequencing (PLATE-Seq) and 1 plate for viability testing in comparison to peripheral Tregs at the peripheral Treg IC20 dose. Seven drugs with significantly greater toxicity to tumor Tregs vs peripheral Tregs were identified (Figure 3C).

Wells of drug-treated Tregs were RNA-Sequenced and each normalized with viperSignature against the internal DMSO-control wells on the same PLATE. VIPER was run on the normalized gene expression using the T-cell ARACNe network inferred from sorted bulk-RNA-Sequencing clinical data. Drugs were ranked on their overall inversion across patients of the 17-gene Master Regulator signature previously identified and validated by CRISPR (Figure S3B), as well as on their patient-by-patient inversion of Tumor-Treg vs Peripheral-Treg protein activity signature by OncoTreat (Figure 3E).

### Tumor-Growth Screens

We assessed tumor growth first in response to treatment with floxuridine, triapine, and gemcitabine relative to untreated control, with or without anti-PD-1 immunotherapy (Fig 4A-C). 10 C57BL/6J mice per treatment arm were implanted with subcutaneous MC38 tumor cells. Treatment was initiated after 12 days of initial tumor growth, at which point mice were monitored for tumor volume until exceeding 1000mm^3 or ulceration exceeding a diameter of 5mm. Gemcitabine was administered IP on days 12, 15, and 18 at 12 mg/kg, or 1/10^th^ of the lowest conventional clinical-equivalent dose in mice (120 mg/kg). Floxuridine and triapine were IP daily from day 12-18 at 1mg/kg and 5mg/kg, respectively, also reflecting 1/10^th^ the standard murine dose. Mice receiving anti-PD-1 were administered anti-PD-1 IP on days 12, 15, and 18. Treatment response outcomes were assessed by cox proportional hazards model (Figure 5C), Kaplan-Meier curve (Figure 5B), and computation of mean tumor growth slope over time (Figure S4). Tumor growth curves display the average tumor volume over time with standard deviation represented by error bars or shading, with each average growth curve terminating when the first animal in each treatment condition reaches end stage (1,000 mm^3^). By all criteria, gemcitabine was the only treatment found to significantly inhibit tumor growth, alone and in combination with anti-PD-1.

To further assess the doses at which gemcitabine inhibits tumor growth and the immune-mediated effects of gemcitabine, we performed parallel dose titrations of Gem in immune-competent C57BL/6J mice and immune-deficient NSG (NOD.Cg-*Prkdc^scid^ Il2rg^tm1Wjl^*/SzJ) mice, administering doses ranging from 0.12mg/kg up to 120mg/kg as shown in Figures 4D-E. At least 5 mice were treated per treatment arm. Doses were administered IP on days 12, 15, and 18, and treatment response was assessed by and Kaplan-Meier test (Figure 4E).

Finally, tumor growth was assessed in single-gene TRPS1 CRISPRko generated by the CHIME protocol described above, compared to transduction by CHIME with a non-targeting scramble control guide. These cohorts included 6 TRPS-KO mice and 5 Scramble-control mice. For these mice, we pooled two guides targeting TRPS1 and two non-targeting guides with approx. MOI 50 based on 293T cell line tittering. These guide sequences were:

**TRPS1_1:** AGAGGGGCAGACATCCTACG

**TRPS1_2:** AGCATCGGATGTCAAACAGG

**Non-targeting guide 1:** GCGAGGTATTCGGCTCCGCG

**Non-targeting guide 2:** GCTTTCACGGAGGTTCGACG

Following immune reconstitution, mice were initially implanted with subcutaneous MC38 tumor, which spontaneously regressed in both arms following initial tumor growth for two weeks post-implantation. This was assumed to be driven by baseline immune activation caused by irradiation and stem cell transplant coupled with the higher MOI used and immune sensitivity of early post-implantation MC38 tumors. Subsequently, these mice were implanted on the opposite flank with subcutaneous MCA205, a more aggressive and immune-resistant fibrosarcoma cell line. Tumor volume was assessed every 48 hours following day 7 post-implantation, such that tumor volumes in TRPS1 mice were determined to be significantly lower than scramble controls by day 13 (p < 0.05). Treatment response was assessed by Kaplan-Meier test.

### Single-Cell RNA-Seq Profiling of Gemcitabine Effect on TI-Tregs

To test the hypothesis that low-dose Gem modulates TI-Tregs, we performed single cell RNA sequencing of MC38 tumor- and spleen-derived Tregs 24 hours after exposure to a single dose of 12 mg/kg Gem as well as 24 hours after vehicle control. For this study, we implanted *FoxP3^Yfp-Cre^* mice with MC38 to facilitate flow-sorting of TCR-β^+^ CD4^+^ FoxP3^+^ Tregs from tumor and spleen specifically by the YFP marker. Tissue was harvested at day 14 post tumor-implantation, and fresh tissue was minced to 2-4 mm sized pieces in a 6-cm dish and subsequently digested to single cell suspension using Multi Tissue Mouse Tumor Dissociation Kit 1 (Miltenyi Biotec) and a gentleMACS OctoDissociator (Miltenyi Biotec) according to the manufacturer’s instructions.

Dissociated cells were flow-sorted for YFP^+^ Tregs and processed for single-cell gene expression capture (scRNASeq) using the 10X Chromium 3’ Library and Gel Bead Kit (10x Genomics), following the manufacturer’s user guide at the Columbia University Genome Center. After GelBead in-Emulsion reverse transcription (GEM-RT) reaction, 12-15 cycles of polymerase chain reaction (PCR) amplification were performed to obtain cDNAs used for RNA-seq library generation. Libraries were prepared following the manufacturer’s user guide and sequenced on Illumina NovaSeq 6000 Sequencing System. Single-cell RNASeq data were processed with Cell Ranger software at the Columbia University Single Cell Analysis Core. Illumina base call files were converted to FASTQ files with the command “cellranger mkfastq.” Expression data were processed with “cellranger count” on pre-built mouse reference. Cell Ranger performed default filtering for quality control, and produced a barcodes.tsv, genes.tsv, and matrix.mts file containing transcript counts for each cell, such that expression of each gene is in terms of the number of unique molecular identifiers (UMIs) tagged to cDNA molecules corresponding to that gene.

These data were loaded into the R version 3.6.1 programming environment, where the publicly available Seurat package was used to further quality-control filter cells to those with fewer than 25% mitochondrial RNA content, more than 1,000 unique UMI counts, and fewer than 15,000 unique UMI counts. Pooled distribution of UMI counts, unique gene counts, and percentage of mitochondrial DNA after QC-filtering is shown in Figure S5A. Gene Expression UMI count matrix was processed in R using the Seurat SCTransform command followed by Seurat Anchor-Integration. The sample was clustered on gene expression by a Resolution-Optimized Louvain Algorithm (Hao et al., 2021). Protein activity was inferred for all cells by VIPER using the SCTransform gene expression signature and the T-cell ARACNe network derived from sorted T-cell bulk-RNA-Seq. The single-cell data were then re-clustered on VIPER protein activity (Figure S5B). Top 5 most differentially upregulated proteins per cluster were assessed by t-test (Figure S5C). Enrichment of the TI-Treg MRs was assessed by Gene Set Enrichment Analysis (GSEA) on a cell-by-cell basis, with normalized enrichment scores shown in Figure 5E and protein activity of the individual MRs shown in Figure 5D. Cluster frequencies were plotted for each sample (Vehicle-Treated Tumor, Vehicle-Treated Spleen, Gem-Treated Tumor, Gem-Treated Spleen), with pairwise comparisons in frequency assessed by Fisher’s Exact test and cox proportional hazards model (Figure 5G).

---

**Figure S1.**
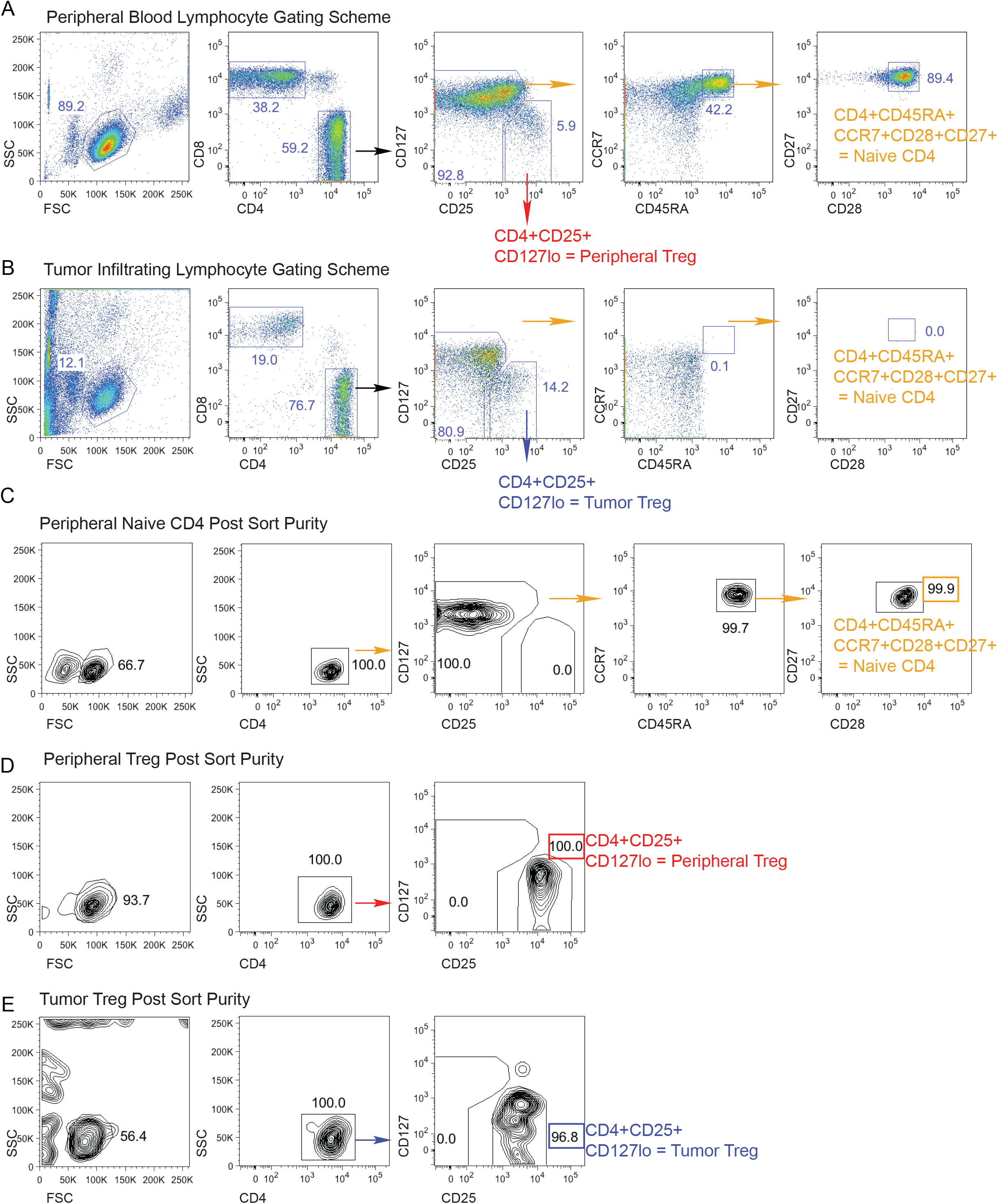
Human T Cell Sorting Schemes and Purity: **(A)** Sorting scheme for flow isolation of peripheral blood CD8, Treg, and naïve CD4 Tconv populations. **(B)** Sorting scheme for flow isolation of tumor infiltrating CD8, Treg, and naïve CD4 Tconv populations. **(C)** Purity of peripheral blood naïve CD4 T cells post-sort. **(D)** Purity of peripheral blood Tregs post-sort. **(E)** Purity of TI-Tregs post-sort.

**Figure S2:**
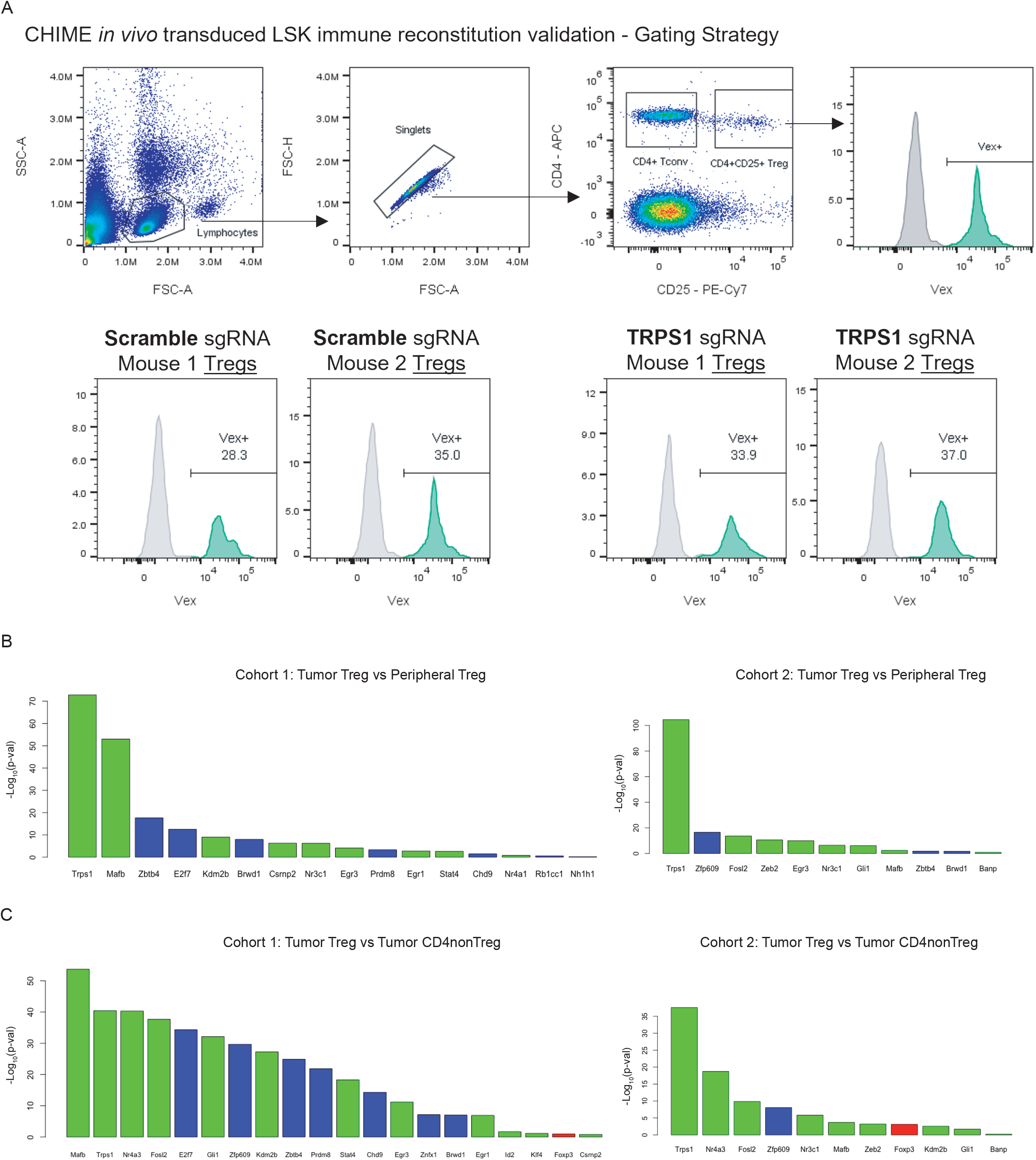
CRISPRko Transduction Efficiency and Individual Cohort Results: **(A)** Flow gating schema and sgRNA transduction efficiency following immune reconstitution of CHIME animals. Vex+ frequencies of peripheral blood Treg populations are shown for two representative mice per condition. **(B)** Barplot of -log10(Bonferroni-Corrected P-values) for genes with statistically significant depletion of targeting gDNAs in Tumor Tregs vs Peripheral Tregs, separately for experimental cohort 1 (left) and experimental cohort 2 (right). **(C)** Barplot of -log10(Bonferroni-Corrected P-values) for genes with statistically significant depletion of targeting gDNAs in Tumor Tregs vs Tumor CD4nonTregs, separately for experimental cohort 1 (left) and experimental cohort 2 (right).

**Figure S3:**
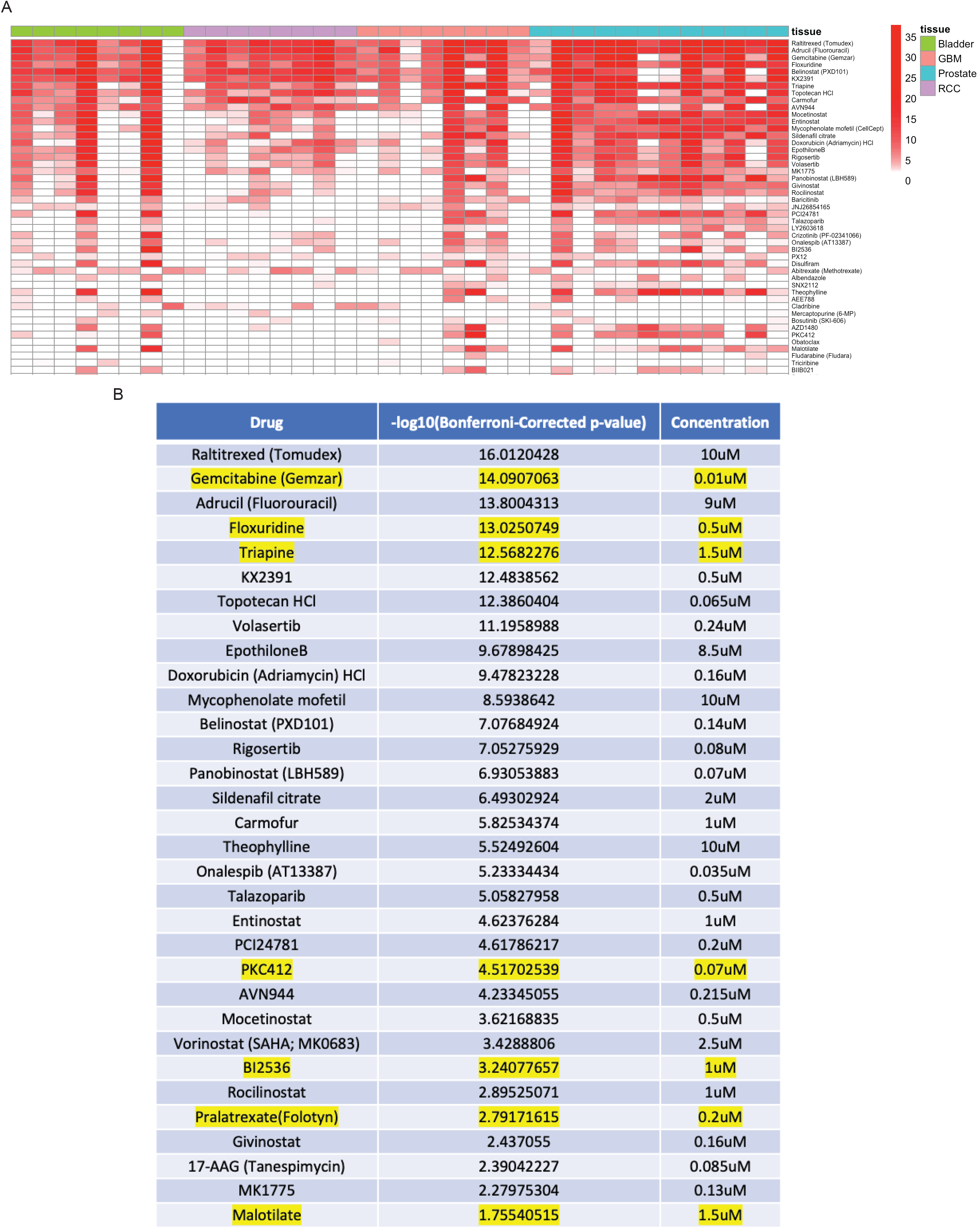
Tumor-Treg OncoTreat Drug Predictions, Expanded List of All Statistically Significant Compounds: **(A)** Patient-by-Patient Drug predictions according to inversion of patient-specific Tumor Treg vs Peripheral Treg protein activity signature by drug-treatment protein activity signature. Each drug predicted to invert Tumor Treg signature with - log10(Bonferroni-Corrected p-value) < 0.01 in a particular patient is colored red. Patients are grouped by tumor type. **(B)** Table of all drugs significantly down-regulating Tumor-Treg MRs identified in Figure 1E, 1F, ordered by p-value. Drugs also identified by growth screen to have differentially higher toxicity in Tumor Tregs vs Peripheral Tregs are highlighted in yellow. All seven of these are identified as statistically significant hits down-regulating Tumor-Treg MRs.

**Figure S4:**
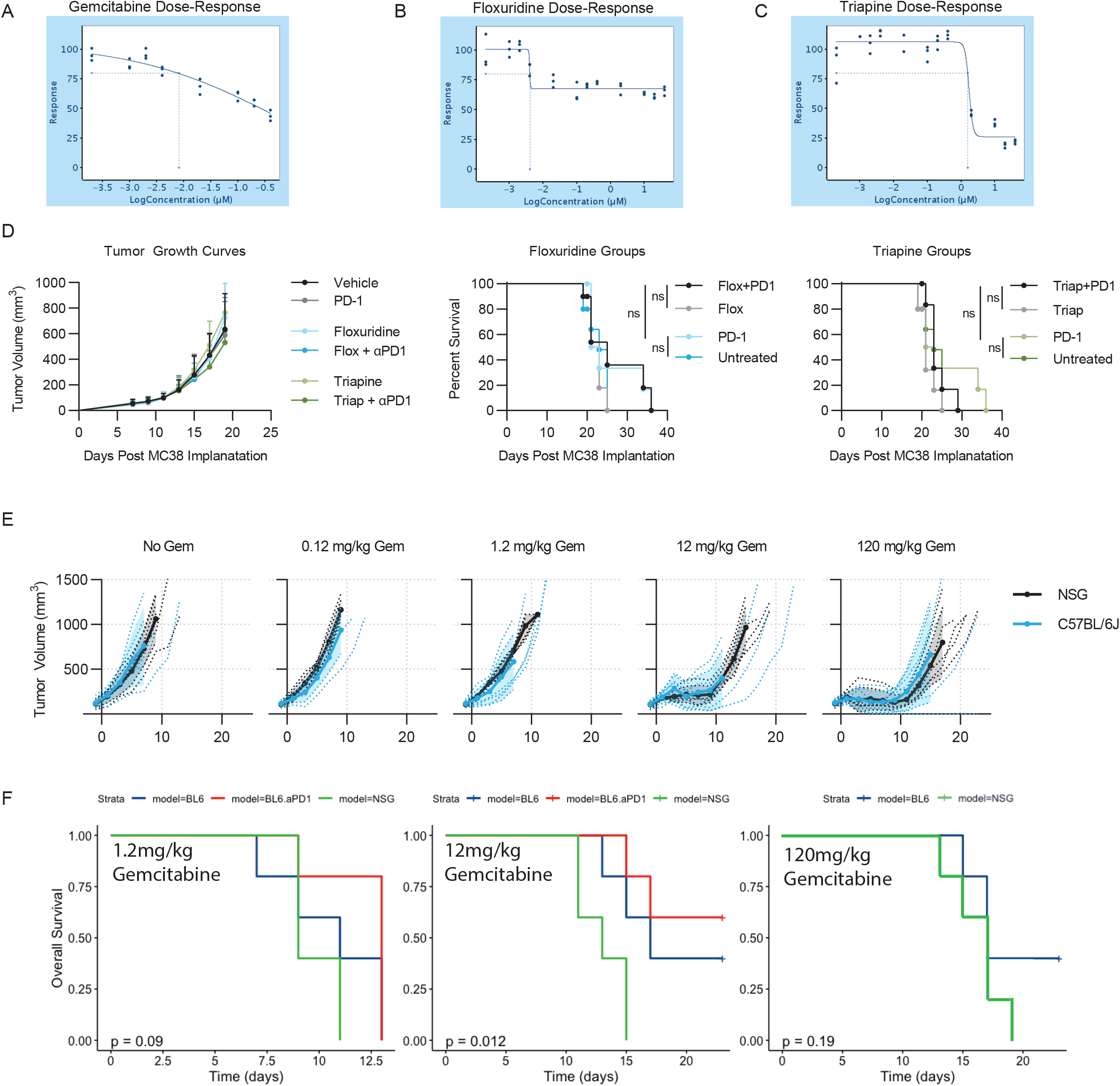
In-vivo Tumor Growth Inhibition by Gemcitabine, Floxuridine, and Triapine: **(A)** Dose-Response titration curve for gemcitabine on ex vivo Treg growth inhibition. **(B)** Dose-Response titration curve for floxuridine on ex vivo Treg growth inhibition. **(C)** Dose-Response titration curve for triapine on ex vivo Treg growth inhibition. **(D)** Tumor growth curves over time for each treatment (vehicle, floxuridine, triapine, anti-PD-1, anti-PD-1+floxuridine, anti-PD- 1+triapine), Kaplan-Meier curves for floxuridine treatment groups and triapine treatment groups. **(E)** Tumor growth curves of gemcitabine titration at 120 mg/kg, 12 mg/kg, 1.2 mg/kg, and 0.12 mg/kg doses in NSG vs C57BL/6J mice. **(F)** Kaplan-Meier curves of gemcitabine dose titration at 1.2mg/kg, 12mg/kg, and 120mg/kg, showing comparison of overall survival time in BL6 immune-competent mice (with or without aPD1) versus NSG immunodeficient mice (p=0.09, p=0.012, and p=0.19, respectively)

**Figure S5:**
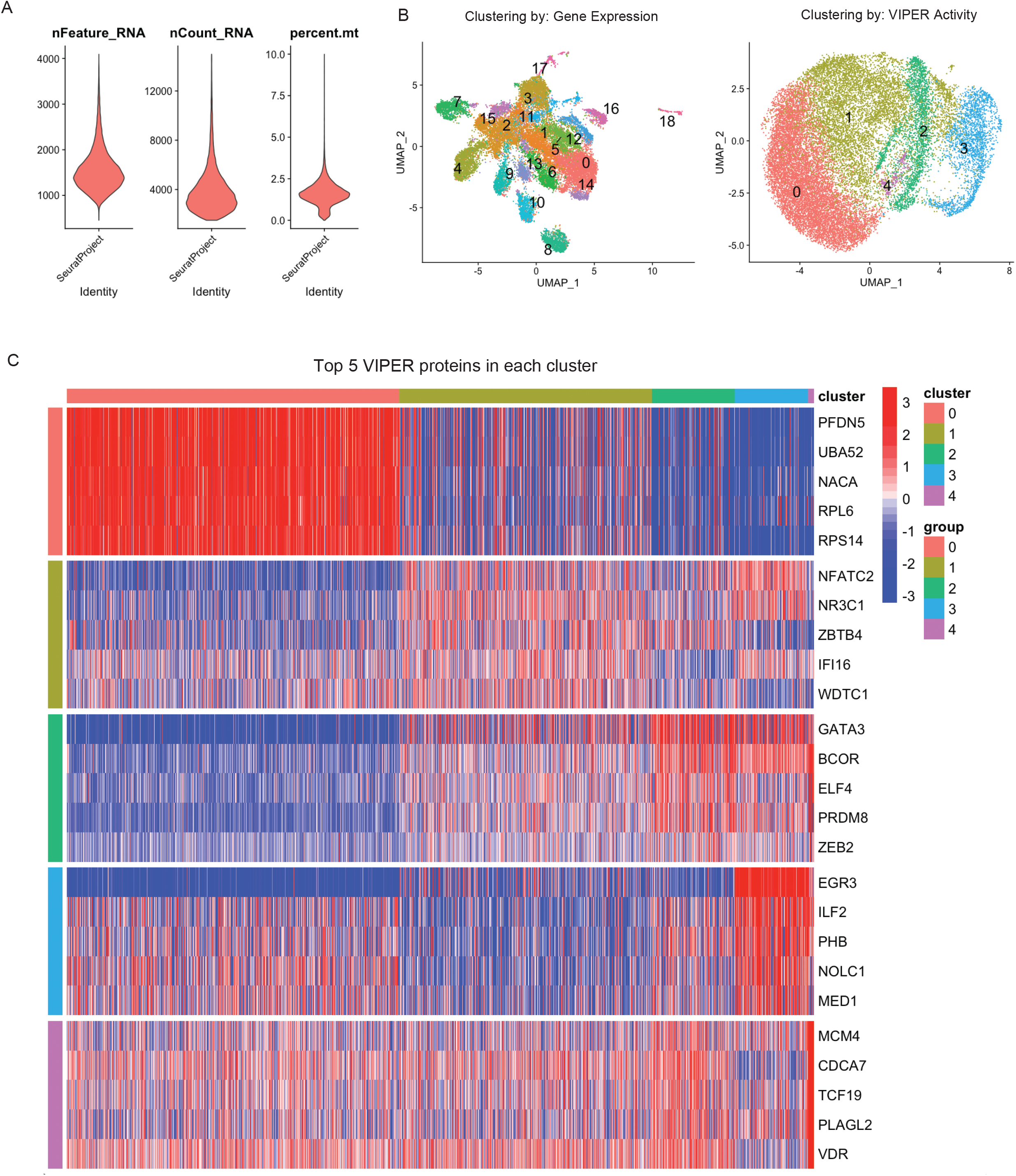
Single-Cell RNA-Seq Characterization of Tumor-Infiltrating and Peripheral Tregs With or Without Low-Dose Gemcitabine Treatment: **(A)** Violin plot of data quality showing distribution of nFeature_RNA (number of unique genes profiled), nCount_RNA (number of unique molecular identifiers profiled), and percent.mt (percentage of mitochondrial transcripts) per cell. **(B)** Clustering of Tregs by Gene Expression (left) and VIPER protein activity inference (right), showing noisiness of clustering by gene expression due to cross-sample batch effects. **(C)** Top5 most differentially upregulated proteins per Treg cluster.

## Author Contributions

A.O. conceived of, coordinated, assisted in performing all experiments, and performed data analysis, and co-wrote the manuscript. C.A. performed and designed tumor growth and tumor-infiltration screens and designed all flow cytometry panels. M.T. optimized and performed all CRISPR transduction protocols. M.M. assisted in hematopoietic stem cell transplantation. C.J, T.N., C.K, M.A., T.B., V.Y., A.D., and M.L. recruited patients and coordinated collection of tumor and peripheral blood specimens for sorted T-cell bulk-RNA-Seq. C.K. ran the high-throughput drug screen PLATE-Seq assay. A.C. and C.D. conceived of the project, advised on overall experimental design and data analysis procedure, and co-wrote the manuscript.

## Acknowledgements

This research was supported by National Institutes of Health (NIH) grants R01 CA127153, 1P50CA58236-15 and P30CA006973, and CUMC institutional funds to Charles Drake, by NIH grants R35CA197745, U01DA217858, S10 OD012351 and S10 OD021764 to Andrea Califano, by NIH grant F30CA260765-01 to Aleksandar Obradovic, and by NIH grants UL1TR001873 and TL1TR001875 to Casey Ager.

